# ZBTB48 is a pioneer factor regulating B-cell-specific CIITA expression

**DOI:** 10.1101/2023.07.25.550481

**Authors:** Grishma Rane, Vivian L. S. Kuan, Suman Wang, Michelle Meng Huang Mok, Vartika Khanchandani, Julia Hansen, Ieva Norvaisaite, Wai Khang Yong, Arne Jahn, Vineeth Mukundan, Yunyu Shi, Motomi Osato, Fudong Li, Dennis Kappei

**Author notes:** Correspondence should be addressed to Dennis Kappei.

## Abstract

CIITA is the master regulator of MHC II gene expression and hence the adaptive immune response. CIITA expression itself is tightly regulated by three cell type-specific promoters, pI, pIII, and by pIV, and can also be induced by IFNγ in non- immune cells. While key regulatory elements have been identified within these promoters, knowledge of transcription factors regulating CIITA is incomplete. Here, we demonstrate that the telomere-binding protein and transcriptional activator ZBTB48 directly binds to both the critical activating elements within CIITA pIII and is essential for its gene expression. ZBTB48 establishes open chromatin at CIITA pIII upstream of activating H3K4me3 modifications both priming CIITA transcription for IFNγ-induction and ensuring constitutive expression in primary murine B cells. Hence, ZBTB48 acts as a molecular on-off-switch for B-cell-specific CIITA expression.

## INTRODUCTION

Major histocompatibility complex class II (MHC II; also known as HLA class II) cell surface protein complexes are essential for the initiation, amplification and regulation of adaptive immune responses and the development of an immunological memory. Precise expression and assembly of MHC II complexes are thus critical for protection against pathogens, curbing tumour growth as well as moderating autoimmune responses^1, 2^. While MHC II is constitutively expressed on professional antigen presenting cells (APCs) such as macrophages, dendritic cells and B cells, it can also be induced by cytokines such as interferon-γ (IFNγ)^3^ in non-APCs^4^.

The co-ordinated expression of all the MHC II genes is regulated by four common cis-acting DNA elements, which are bound by specific factors that collectively form the MHC II enhanceosome. Three ubiquitously expressed direct DNA binding proteins, RFX^5–8^, NF-Y^9, 10^ and CREB^11^ serve as a combined interaction platform for the MHC II master regulator CIITA (also known as MHC2TA)^12^. CIITA functions as an essential, non-DNA binding co-activator and orchestrates chromatin modification and remodeling to activate MHC II transcription. As the critical upstream regulator, CIITA mirrors the MHC II expression pattern with constitutive expression in APCs and inducible expression in non-APCs, e.g. upregulation upon IFNγ treatment via JAK- STAT signalling^13, 14^. Mutations in CIITA and other MHC II enhanceosome genes cause the bare lymphocyte syndrome (BLS), which can also be recapitulated with BLS mouse models, presenting clinical and immunological defects that can be directly attributed to MHC II deficiency^5–8, 15, 16^. In addition, repression of CIITA is a commonly employed mechanism for MHC II downregulation in diseases with inflammatory components^2^ and by pathogens and cancer cells to avoid immune recognition^4, 17^. For instance, CIITA gene fusions that occur frequently in lymphoid cancers concomitantly result in loss of MHC II expression^18^. Overall, CIITA is essential for both constitutive and inducible expression of MHC II as the master regulator of the adaptive immunity gene expression program.

CIITA transcription itself is controlled by a 14kb multi-promoter regulatory region, containing three independent promoter elements in humans, promoter pI, pIII and pIV. Originally, a pII element had also been identified but its expression has not been detected in human tissues^19–21^. PI, pIII and pIV, each transcribe an isoform with a unique first coding exon and are predominantly expressed in a tissue-specific manner^19^. Constitutively elevated levels of pI expression is found in dendritic cells and macrophages^19, 20^, while pIII is the major contributor for constitutive CIITA expression in B cells, human activated T cells and plasmacytoid dendritic cells^19, 20, 22^. Although all three promoters can be induced by IFNγ to varying degrees, pIV is considered as the predominant responder to induction and thus crucial for IFNγ-induced CIITA expression in non-hematopoietic cells^23, 24^. B-cell-specific CIITA expression depends on a 319bp promoter element upstream of the transcription start site (TSS)^25, 26^. Prior *in vivo* genomic footprint analysis of this CIITA pIII core promoter region had identified two sequences, activation response element (ARE) 1 and 2, that are essential for CIITA expression in B cells^22, 25–27^. While putative candidate proteins have been suggested, these studies had largely been limited to EMSA assays probing for supershifts^22, 25^ and a genuine direct binder for the ARE elements has not been conclusively identified.

ZBTB48 (also known as HKR3 or TZAP) was previously reported as a direct telomere binding protein that acts as a negative regulator of telomere length^28–30^. In addition, it also binds to proximal promoters of a defined set of genes and moonlights as a transcriptional activator^28^. Here, we show that ZBTB48 binds directly to the two ARE sites within the core CIITA pIII. We demonstrate that by establishing open chromatin, ZBTB48 regulates inducible CIITA pIII expression and that constitutive B-cell-specific expression is affected in a ZBTB48 knock-out mouse model leading to a reduction in MHC II positive cells in primary B cells. Overall, we establish that ZBTB48 has pioneering activity and acts as a molecular on-/off-switch upstream of activating histone modifications and gene expression activation.

## RESULTS

### ZBTB48 directly binds to two sites within CIITA pIII

In addition to its function as a telomere length regulator, we have previously identified ZBTB48 as a transcriptional activator based on its preferential binding to proximal promoter regions and consequently lower transcript levels of the matched genes in ZBTB48 KO cells^28^. We had originally prioritised ZBTB48 ChIP-seq peaks with concomitant changes in mRNA expression levels comparing both U2OS and HeLa ZBTB48 WT and KO clones. Due to its lack of constitutive expression in non-immune cells, CIITA transcripts were not detected in the RNA-seq data. However, revisiting our ChIP-seq data revealed that ZBTB48 clearly occupies the region corresponding to CIITA pIII (**Figure 1A**). Given the IFNγ-inducible nature of CIITA expression we reasoned that CIITA might represent a previously unappreciated ZBTB48 target gene and we hence decided to evaluate its functional relevance in MHC II-driven immune response.

**Figure 1:**
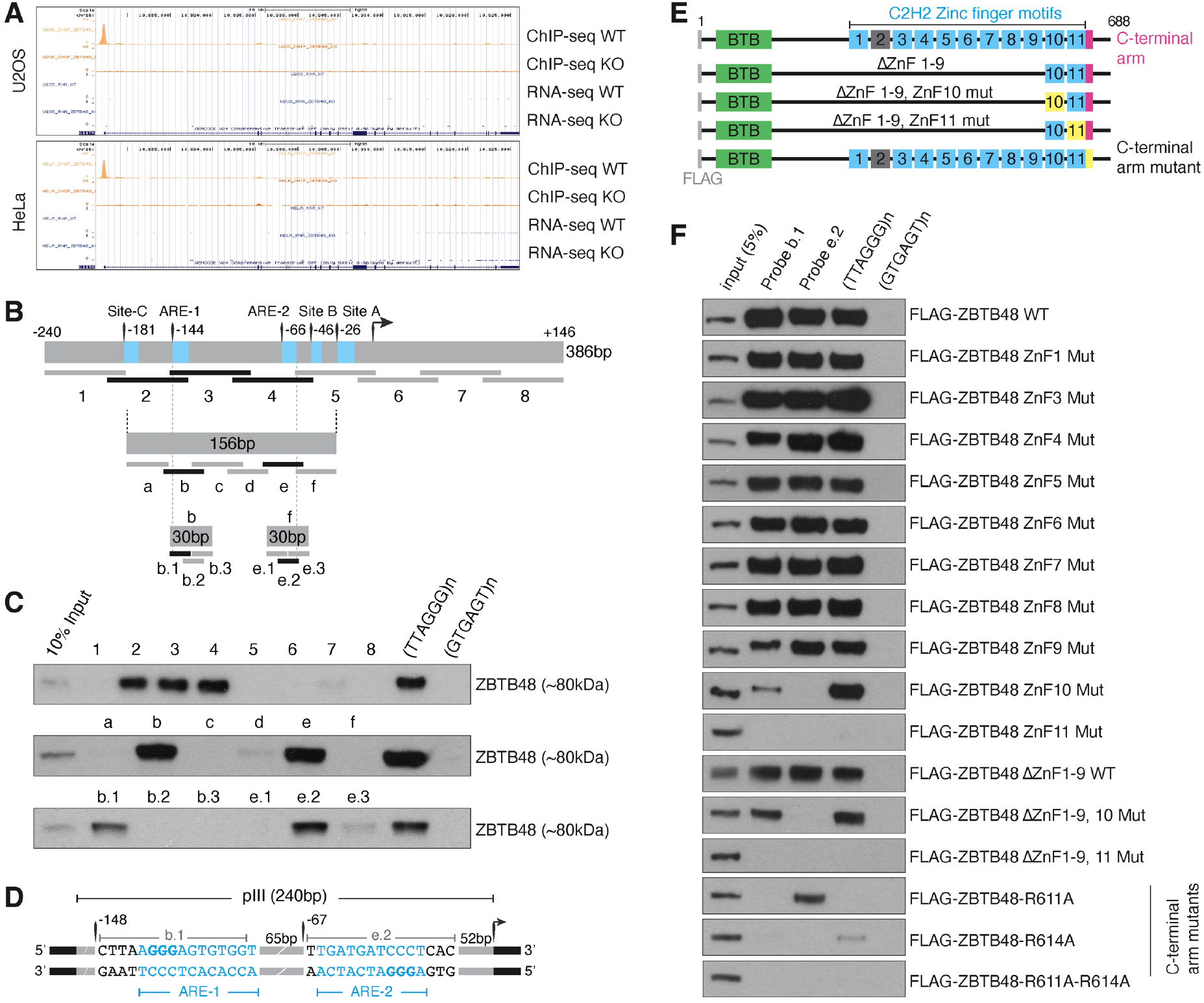
ZBTB48 directly binds to CIITA pIII. (A) ChIP-seq tracks depicting ZBTB48 binding peaks at CIITA pIII in U2OS and HeLa WT cells but not in the KO cells (n=4). No reads are observed in the corresponding RNA-seq tracks in these uninduced cells (n=5; data re-analysed from Jahn *et al*.^28^). (B) Schematic of DNA probe design encompassing the binding peak that was used in the DNA pull-down assay. Transcription start site (TSS) and known regulatory elements are indicated and binding in the pulldowns is denoted by probes in darker shade. (C) Western blot of DNA pull-down assay using U2OS nuclear extracts demonstrates ZBTB48 binding at two sites within pIII. Telomeric (TTAGGG) and scramble control (GTGAGT) sequence are used as positive and negative controls, respectively. (D) Schematic of ZBTB48 binding sites within CIITA pIII with their relative distances to the TSS and their sequence overlap to the originally mapped ARE elements^25^. (E) Schematic of FLAG- tagged ZBTB48, a 688-amino acid protein, containing N-terminal BTB domain (green), 11 zinc fingers (ZnFs) and a C-terminal arm (pink). The functional ZnFs are in blue and degenerate ZnF2 is in grey. Deletion constructs lacking ZnF1-9 with and without points mutations in either ZnF10 or ZnF11 and WT ZBTB48 with mutations in C- terminal arm are depicted. (F) DNA pull-downs with CIITA pIII probes b.1, e.2, telomeric (TTAGGG) and control sequence (GTGAGT) for FLAG-ZBTB48 WT, ZnF mutants and C-terminal arm mutants according to Fig. 1D.

To identify the exact binding site(s) within the ChIP-seq peak, we performed *in vitro* reconstitution DNA pull-downs using nuclear protein extracts from U2OS cells. Eight overlapping oligo probes straddling the ChIP-seq binding peak were designed as shown in **Figure 1B**. As a positive control we used a telomeric (TTAGGG) DNA probe, for which we had previously demonstrated ZBTB48 binding^28^. Here, three probes (2, 3 and 4) showed clear enrichment of ZBTB48, at par with the enrichment observed with the telomeric DNA control, suggesting multiple binding sites within the CIITA pIII region (**Figure 1B and C**). Therefore, using the sequence covered by probes 2-4 as a template, we generated six smaller, 30bp probes (a-f) and observed ZBTB48 enrichment on two discrete probes, b and e. Subsequently, to narrow down the exact binding sites, we generated three overlapping 15bp probes that each span the sequences of the b and e probes. Specific binding of ZBTB48 was identified to probes b.1 and e.2, that localize to -133 to -148bp and -52 to -67bp of the CIITA pIII, respectively, demonstrating that ZBTB48 binds at two different sites within the promoter. Intriguingly, these binding sites correspond to both ARE-1 and ARE-2 (**Figure 1D**), the critical regulatory elements of CIITA pIII expression.

### ZBTB48 interacts with CIITA pIII via its ZnF10, ZnF11 and C-terminal arm

We have previously demonstrated that among the 10 functional zinc fingers in ZBTB48 (ZnF2 is degenerate), the 11^th^ zinc finger (ZnF11) in combination with a short C-terminal arm are necessary and sufficient for binding to telomeric DNA^28, 31^. To test whether binding at b.1 and e.2 is also mediated by ZnF11, we expressed point mutants of FLAG-ZBTB48 in U2OS cells by individually exchanging the first histidine within each Cis_2_His_2_ ZnF motif to alanine and performed DNA pull-downs using the b.1 and e.2 probes (**Figure 1E**). While FLAG-ZBTB48 WT and point mutants of ZnF1-9 efficiently bound to both b.1 and e.2, mutation of ZnF11 (ZnF11 mut) resulted in complete loss of its binding ability (**Figure 1F**) in parallel to the expected loss in TTAGGG binding. In contrast, the ZnF10 mutant (ZnF10 mut) is capable of binding to telomeric DNA, but its binding to b.1 was strongly reduced and completely lost on e.2. We further confirmed the requirement of both ZnF10 and ZnF11 by using deletion constructs that lack ZnF1-9 and carry the same point mutations in either ZnF10 or ZnF11 (**Figure 1E**). Indeed, FLAG-ZBTB48 ΔZnF1-9 was enriched on both the probes, but a mutation in ZnF10 led to reduced enrichment on probe b.1 and loss of enrichment on e.2, while a mutation in ZnF11 led to a complete loss of binding ability on all probes (**Figure 1F**). In addition, results from a previous co-crystal structure with telomeric DNA demonstrated that ZnF11 recognizes the G_4_G_5_G_6_ while two amino acid residues from the adjacent C-terminal arm, R611 and R614, are responsible for the recognition of T_2_A_3_^31^. To test whether the adjacent C-terminal arm is also required for CIITA pIII binding we tested R611A and R614A mutants in our pull-down assay (**Figure 1E**). While the R611A mutant retained some enrichment on e.2 and R614A on telomeric DNA, neither mutant showed any binding to b.1. Furthermore, the R611A/R614A double mutant was incapable of binding to any of the 3 sequences (**Figure 1F**). These results demonstrate that ZBTB48 binds at two distinct sites within proximal CIITA pIII and requires ZnF10, ZnF11 and the adjacent C-terminal arm.

To further confirm the involvement of ZnF10 in binding to the CIITA promoter we generated a co-crystal structure of the e.2 probe sequence with a construct containing ZnF10-11 as well as the C-terminal arm^31^ (**Figure 2A**). Similar to telomeric DNA, base-specific interactions are conferred by ZnF11 in the major groove as well as by the C-terminal arm that turns back and lies across the phosphate backbone to fit in the minor groove of the double-stranded e.2 DNA (**Figure 2B, Table S1**). Here, the RxxHxxR motif of ZnF11 recognizes G_10_G_11_G_12_ with Arg589, His592 and Arg595 forming a bidentate hydrogen bond interaction with G_12_, G_11_ and G_10_, respectively. This is equivalent to the interaction of both ZBTB48 and ZNF524 with telomeric repeats^32^ and ZBTB10 with the telomeric variant repeat TTGGGG^33^ (**Figure 2C**). However, in contrast to the co-crystal structure with telomeric DNA^31^, ZnF10 does not lie outside of the CIITA e.2 DNA duplex but rather interacts with the phosphate backbone along the major groove (**Figure 2B, Supplemental Figure S1**). In the C-terminal arm, of the two arginine residues that conferred base-specific contacts with T_2_A_3,_ R614 forms hydrogen bonds with G_12_ and on the complementary strand with C_8_T_9_ while R611 interacts with the phosphate backbone (**Figure 2A and 2D**). In addition to establishing the molecular details underlying the interaction of the ZBTB48 ZnF10, ZnF11 and the C-terminal arm with the CIITA pIII sequence, these results establish potential separation of function mutations for ZBTB48’s role at telomeres versus as a transcriptional activator – at least for binding to the CIITA pIII.

**Figure 2:**
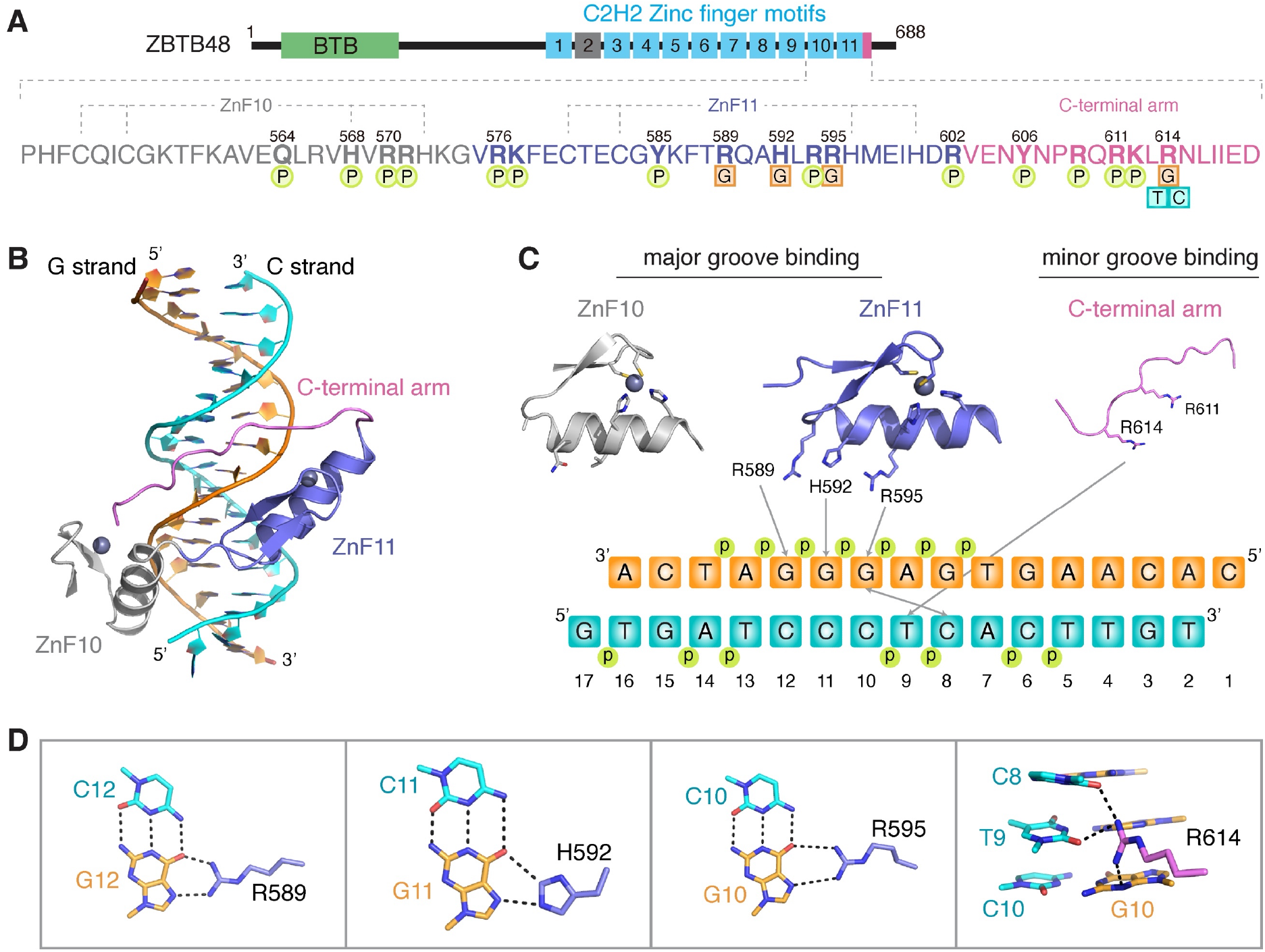
ZBTB48 binding to CIITA pIII requires ZnF10 and ZnF11. (A) Schematic representation of the ZBTB48 protein structure and a zoom in on the expression construct containing ZnF10, ZnF11 and the C-terminal arm. Base-specific interactions (squares) and interactions with the DNA phosphate backbone (circles) are indicated. (B) Overall co-crystal structure of the essential ZBTB48 DNA-binding domains with the e.2 binding site within CIITA pIII. (C) Base-specific interactions depicted for each of the three ZBTB48 domains involved in binding to CIITA pIII. Binding of ZnF11 to the GGG sequence involves its RxxHxxR motif. (D) Details of base-specific interactions for ZnF11 and the C-terminal arm. Hydrogen bonds are depicted as dashed lines.

### ZBTB48 mediates IFNγ-induced expression of CIITA

ARE-1 and ARE-2 are essential elements in CIITA pIII, governing CIITA expression in B cells^25^. The overlap of ZBTB48 binding sites with these elements suggests a regulatory role of ZBTB48 in CIITA pIII expression. To investigate whether CIITA expression is affected by the loss of ZBTB48, we compared CIITA transcript levels in each five WT and ZBTB48 KO U2OS clones as biological replicates upon IFNγ treatment. Using primers that detect all CIITA isoforms (pan-CIITA), as expected we observed only very low baseline expression of CIITA in untreated U2OS cells (**Figure 3A**). In contrast, IFNγ stimulation resulted in a strong, >1000-fold induction of CIITA in WT cells with a 34-fold higher expression than in the ZBTB48 KO clones. Since ZBTB48 binds at pIII of CIITA, we further assessed whether the difference in pan- CIITA expression specifically arises from lower expression of pIII. Although IFNγ treatment induced expression of all three CIITA promoters, no significant difference in pI expression was observed between WT and ZBTB48 KO clones and, overall, pI induction was modest. pIII was the most strongly induced isoform and its expression was 154-fold higher in WT clones compared to ZBTB48 KO clones (**Figure 3A**). Similarly, transcript levels of pIV, which resides downstream of pIII, were also on average 17-fold higher in the WT clones. The loss of pIII expression was further recapitulated by RNA-seq data from the same five WT and KO clones, where the enrichment of transcripts from the first exon that is unique to pIII is absent in the KO cells after IFNγ stimulation (**Figure 3B**). Overall CIITA transcript levels were abundant in the WT clones with only minimal levels detected in the KO clones, clearly demonstrating that ZBTB48 is critical for IFNγ-induced CIITA pIII expression.

**Figure 3:**
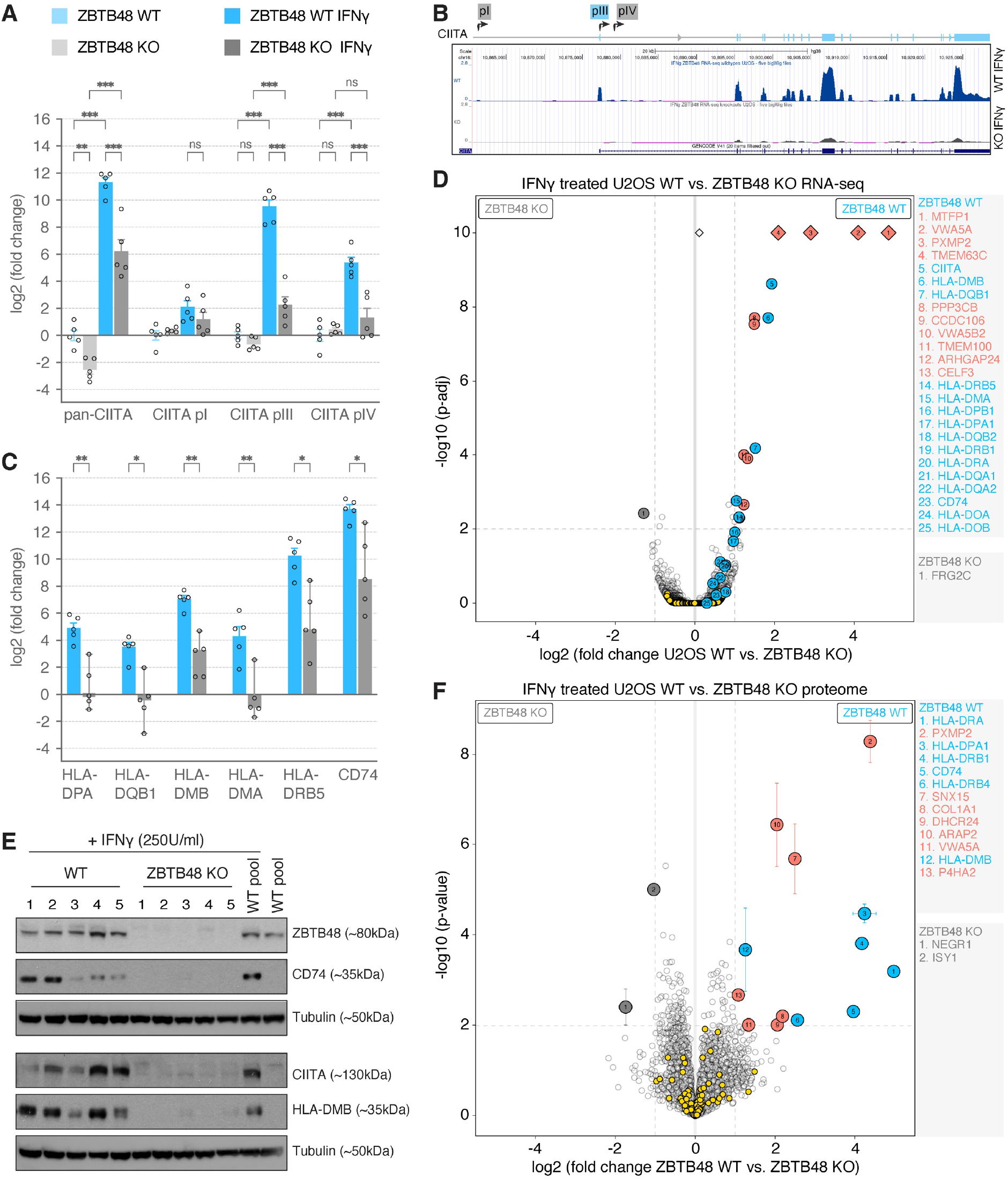
ZBTB48 is required for IFNγ-induced expression of CIITA pIII. (A) Relative mRNA expression of total (pan-)CIITA and the three promoter-specific transcripts in each five independent U2OS WT and ZBTB48 KO clones treated with or without 250 U/ml IFNγ for 24 hours. Data represents mean ± SEM. The log_2_ fold change is calculated relative to the average of five untreated WT clones. p-values were calculated by 2-way ANOVA (n = 5). (B) RNA-seq tracks at the CIITA gene in IFNγ-induced U2OS clones of five WT and ZBTB48 KO clones (n = 5). The positions of the three promoters are indicated in the schematic above. (C) mRNA expression changes of HLA genes in the five U2OS WT and ZBTB48 KO clones treated with 250 U/ml IFNγ for 24 hours. Data represents mean ± SEM. The log_2_ fold change is calculated relative to the average of five untreated WT clones. p-values were calculated by multiple t-test controlling the FDR by two-stage step-up method of Benjamini, Krieger and Yekutieli to correct for multiple comparisons (n = 5). (D) Differential expression analysis of the RNA-seq quantification, comparing each five U2OS WT and ZBTB48 KO clones for U2OS. (E) Western blot for CIITA, HLA-DMB and CD74 protein expression in five U2OS ZBTB48 KO clones compared to five WT clones each when treated with 250 U/ml IFNγ for 48 hours. The parental U2OS WT pool treated with or without IFNγ treatment are included for reference. (F) Protein expression analysis comparing five U2OS WT and ZBTB48 KO clones treated with IFNγ (250 U/ml, 48 hours) by label-free quantitative mass spectrometry. Two- dimensional error bars represent the standard deviation based on iterative imputation cycles during the label-free analysis to substitute missing values (e.g. no detection in the KO clones). For volcano plots in D and F, specifically enriched hits are distinguished from background proteins by a two-dimensional cut-off of >2-fold enrichment and p<0.01. The hits belonging to the CIITA-MHCII family are shown in blue and the rest of the IFNγ response genes (ISGs) in yellow, ZBTB48 targets independent of IFNγ in salmon and hits enriched in KO clones are shown in grey.

To validate the downstream consequence of reduced CIITA expression in ZBTB48 KO cells, we analysed transcript levels of HLA-DPA, -DQB1, -DMB, -DMA, -DRB5 and CD74 as representatives for MHC II expression. Again, upon IFNγ stimulation, U2OS WT clones expressed 15- to 29-fold higher levels of the HLA genes compared to ZBTB48 KO clones (**Figure 3C**). To comprehensively interrogate the effect of ZBTB48 on transcription upon IFNγ stimulation, we globally analysed expression differences in our RNA-seq data from each five U2OS WT and ZBTB48 KO clones as biological replicates treated either with or without IFNγ. Overall, a strong induction of interferon-stimulated genes (ISGs) was observed in both WT and ZBTB48 KO clones upon IFNγ stimulation (**Supplemental Figure S2A and 2B, Table S2**). In agreement with our qPCR data, when comparing the WT and KO clones upon IFNγ treatment, expression levels of CIITA, HLA-DQB1, -DMB and -DMA were significantly higher in the WT clones based on cut-offs of fold change >2 and adjusted p-value < 0.01 (**Figure 3D**). The other MHC II genes were detected but did not achieve significance due to the stringent cut-offs, but overall the expression of the entire gene family was elevated in the WT clones (**Supplemental Figure S2D**). Consistent with our previous data^28^, the expression of MTFP1, VWA5A, and PXMP2, were also decreased in ZBTB48 KO clones both in IFNγ treated and untreated samples (**Figure 3D, Supplemental Figure S2C**). Strikingly, beyond MHC II related transcripts, no other ISGs were differentially expressed upon IFNγ stimulation between ZBTB48 WT and KO cells (**Figure 3D**). These results suggest that the impact of ZBTB48 in response to IFNγ stimulation is exclusive to CIITA expression and that ZBTB48 itself does not respond to IFNγ.

We further validated the lower expression of CIITA and its target genes in IFNγ-treated U2OS ZBTB48 KO clones at protein level by Western blot. In agreement with the qPCR data, CIITA protein levels were low to undetectable across the five ZBTB48 KO clones. Consequently, HLA-DMB and CD74 protein levels were also either below the limit of detection in KO clones or barely detectable and strongly reduced compared to any of the WT clones (**Figure 3E**). To further validate this, we measured global protein expression using label-free quantitative mass spectrometry analysis in five U2OS WT and ZBTB48 KO clones treated with or without IFNγ. While the proteomes are not as comprehensive as the RNA-seq data and did not cover all gene family members, we detected HLA-DRA, -DRB1, -DPA1, -DRB4, DMB and CD74 as members of the CIITA- MHC II axis. As for the RNA-seq data, ISGs were induced in both WT and KO clones upon IFNγ treatment, but again only the MHC II proteins showed significantly higher expression in the WT clones compared to the ZBTB48 KO clones (**Figure 3F, Supplemental Figure S3A-D, Table S3**). Once more, the changes observed between WT and KO cells remained limited to factors that were previously described to differ independent of IFNγ-induction and the CIITA-MHC II axis. In sum, the combined transcriptomic and proteomic profiling validates that in the cascade of cellular IFNγ response, ZBTB48 specifically enables induction of CIITA expression – primarily its pIII transcript –, which in return drives MHC II expression.

### ZBTB48 loss results in diminished MHC II expression *in vivo*

CIITA pIII is conserved in mice including the triple G motifs that are recognised by ZBTB48 through base-specific contacts within the b.1 and e.2 sequences (**Figure 4A**), suggesting that ZBTB48 might be responsible for a similar regulatory mechanism in mice. To test the *in vivo* effects of loss of ZBTB48 on MHC II, we established a ZBTB48 knock-out mouse model in the C57BL/6NTac background. To this end, we injected Cas9 into single-cell stage mouse embryos together with two sgRNAs flanking ZBTB48 exon 2 (which contains the start codon) and a single-stranded oligo DNA nucleotide (ssODN) to guide homology directed repair (**Figure 4B**). The deletion was confirmed by a shorter PCR product as seen in **Figure 4C** and by sequencing (**Supplemental Figure S4A**).

**Figure 4:**
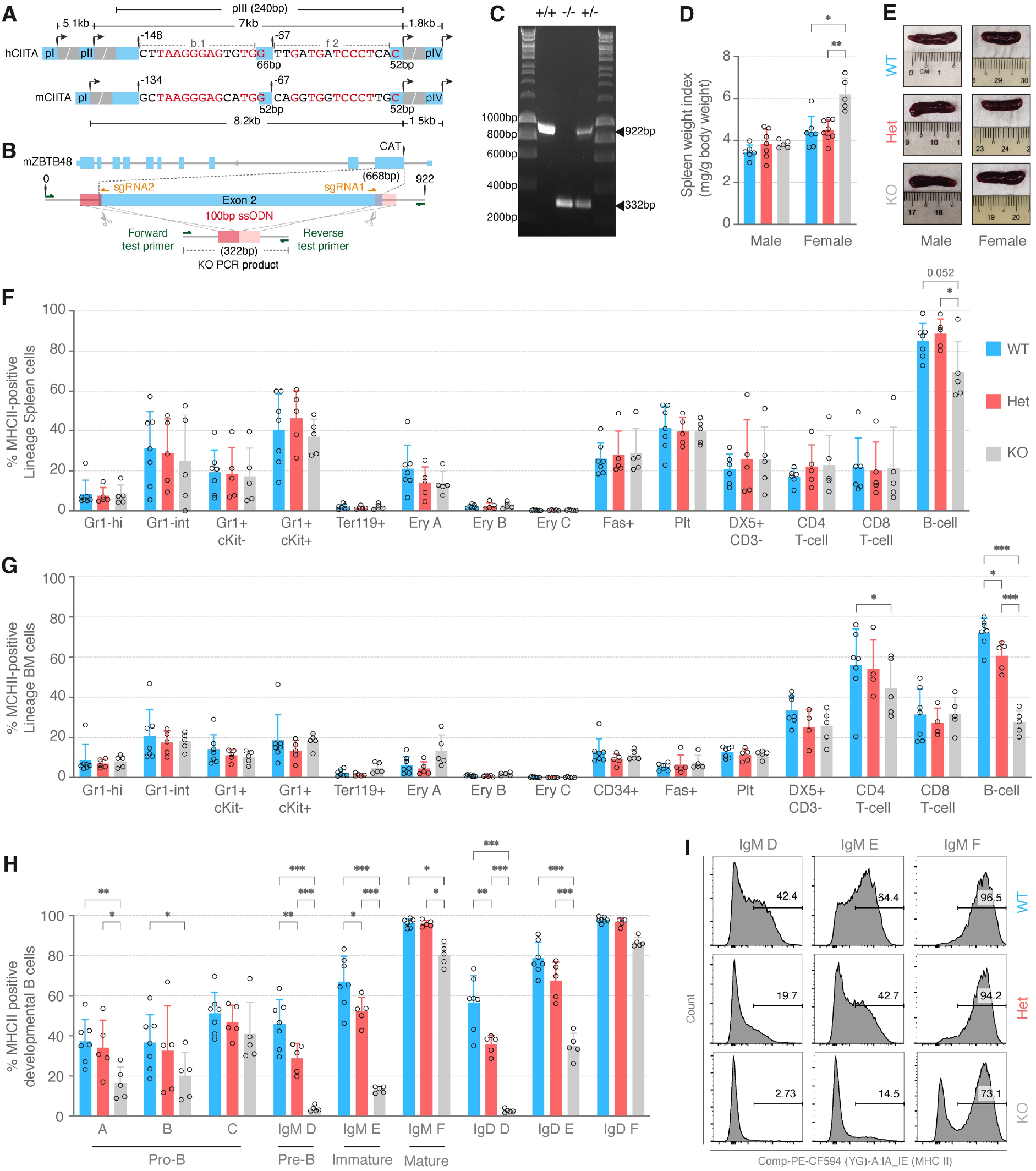
Loss of ZBTB48 diminishes B cell specific MHC II expression *in vivo*. (A) Schematic of the CIITA promoter locus depicting all four promoters in human and three in mice. The sequence conservation between human and mice pIII of b.1 and f.2 is indicated. (B) Schematic overview of sgRNAs and ssODN for CRISPR/Cas9 mediated deletion at exon 2 of the mZBTB48 locus. The ssODN containing 50bp homology arms flanking the cut sites is depicted. The primers (green) used for genotype confirmation flanking the ssODN and the resulting PCR product size is indicated. (C) Mouse genotyping results using primers indicated in (B) and DNA obtained from tail clippings. PCR products from Zbtb48^-/-^ (KO) and Zbtb48^+/-^ (Het) mice were separated on a 1% agarose gel to differentiate between WT (922bp) and KO (322bp). (D) Spleen weight index (weight of spleen in mg / body weight in g) calculated separately for males and females across the three genotypes for 11-14 weeks old mice. Data represents mean ± SD. p-values were calculated by Mann-Whitney test (male: WT n = 6, Het n = 7, KO n = 5; female: WT n = 7, Het n = 8, KO n = 5). (E) Representative spleen images comparing spleen size across the three genotypes in males and females. (F-H) Percentage of MHC II positive lineage specific cells in spleen (F), bone marrow (BM, G) and developmental B cells (H) in WT (n = 7), Het (n = 5) and KO (n = 5) mice. Data represents mean ± SD. p-values were calculated by 2-way ANOVA with Sidak correction for multiple comparisons. The developmental stage of the B cells in H are indicated below. (I) Representative flow cytometry analysis of MHC II positive cells in D (pre-B), E (immature) and F (mature) subpopulations from WT, Het and KO littermates when gated by IgM.

ZBTB48^-/-^ (KO) and ZBTB48^+/-^ (Het) mice were born at Mendelian ratio’s and did not display any obvious growth defects compared to their wild-type (WT) littermates (**Supplemental Figure S4B**). However, female but not male KO mice presented with a mild splenomegaly in 11-14 week old animals (**Figure 4D and 4E**). In contrast, total body weight was unaffected for both males and females across genotypes (**Supplemental Figure S4C**). Given that spleen is a B cell rich organ, we next interrogated molecular phenotypes by assessing MHC II levels. Lineage-specific cells were isolated from peripheral blood, thymus, spleen as well as bone marrow (BM) and their MHC II expression was quantified by fluorescence activated cell sorting (FACS) in the ZBTB48 WT, Het and KO mice. While the relative abundance of the different blood lineages and cell types remained unchanged in both spleen and bone-marrow (**Supplemental Figure S3D and 3F**), the percentage of MHC II positive B cells was reduced in both compartments. Splenic B cells presented with a moderate loss in MHC II positive cells in the KO (69%) mice compared to both WT (85%) and Het (89%) animals (**Figure 4F**). This loss was more pronounced in the BM with approximately three times less MHC II positive B-cells in the ZBTB48 KO (28%) mice as compared to that in WT (72%) mice (**Figure 4G**). In addition, in BM cells a dose-dependent effect was detectable with a significant, intermediary reduction of MHC II-positive B-cells in the Het animals. Given the more pronounced reduction in the BM but a milder reduction of MHC II-positive B-cells in the peripheral blood – 90% in the KO vs 98% in the WT mice (**Supplemental Figure S3F and 3G**) –, we wondered whether the MHC II loss originates during B cell development in the bone marrow. The hematopoietic stem and progenitor cells (HSPCs) in the BM were comparable across the three genotypes and did not display any significant change in the number of the relatively moderate levels of MHC II-positive cells (**Supplemental Figure S5A and 5B**). In striking contrast, while again the relative abundance of cell populations belonging to different B-cell differentiation states did not differ across genotypes (**Supplemental Figure S5C**), developmental B-cell populations, but not thymic developmental T-cell populations (**Supplemental Figure S5D and 5E**), were depleted for MHC II-positive cells. In the early pro-B stage A, the number of MHC II-positive B cells reduced by more than half in the ZBTB48 KO (17%) mice as compared to the WT (37%) and Het (34%) littermates (**Figure 4H**). A similar reduction was seen in pro- B stage B cells while pro-B stage C cells did not present a significant reduction in MHC II levels. Remarkably, the loss of MHC II-positive cells peaked in the pre-B (stage D) subpopulation, where MHC II-positive cells were barely detected in the KO mice (3.7%), compared to 46% and 28.8% MHC II-positive cells in the WT and Het mice, respectively (**Figure 4H**). As the cells developed further, this difference tapered off, with on average five times more MHC II-positive cells in the WT (69%) and four times more in Het (52%) as compared to ZBTB48 KO (13%) mice in the immature (E) subpopulation (Figure 4H). Finally, mature B cells (F subpopulation) in WT animals are nearly all MHC II-positive while ZBTB48 KO mice had ∼20% less MHC II-positive cells, similar to the frequency observed in splenic B cells. We confirmed the results for the pre- to mature B cell subpopulations (stages D-F) by gating with both IgM- and IgD-specific antibodies (**Figure 4I, Supplemental Figure S5F**). In sum, we observed a ZBTB48 dose-dependent reduction in the BM-derived MHC II expressing pro- and pre-B cells, likely due to transcriptional silencing of CIITA pIII. Overall, these data clearly illustrate that ZBTB48 is a critical regulator of B cell-specific MHC II expression *in vivo*.

**Figure 5:**
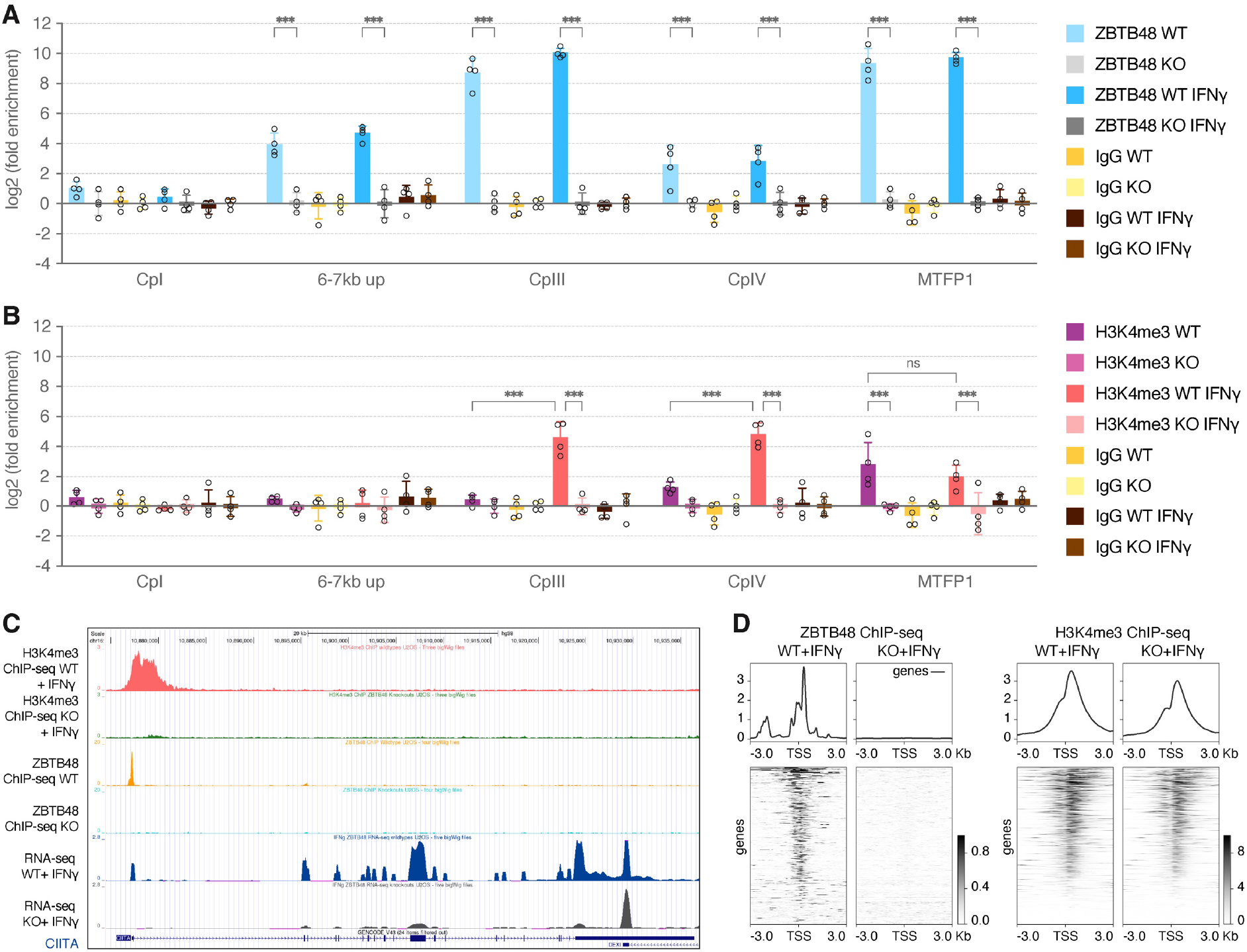
ZBTB48 binding is required for H3K4me3 deposition at CIITA pIII. (A) ChIP reactions from four independent U2OS WT and ZBTB48 KO clones analysed by qPCR for CIITA pI, pII (6-7kb up), pIII, pIV. MTFP1 promoter is used as a positive control. The data was normalised to IgG and gene desert region and fold-change was calculated relative to the average of WT clones. Data represents mean ± SD. p-values were calculated by 2-way ANOVA with Sidak correction for multiple comparisons (n = 4). (B) H3K4me3 ChIP reactions from four independent U2OS WT and ZBTB48 KO clones analysed by qPCR as in (A). (C) ChIP-seq tracks depicting ZBTB48 and H3K4me3 binding peaks at CIITA pIII in U2OS WT cells but not in the KO cells. The H3K4me3 ChIP-seq tracks are cumulative of three IFNγ-induced WT and ZBTB48 KO clones. RNA-seq tracks at CIITA in IFNγ-induced five WT and ZBTB48 KO U2OS clones. The ZBTB48 ChIP-seq tracks are the same as described in Figure 1A. (D) ChIP-seq heatmap at the top 500 ZBTB48 binding sites^28^ (ranked by the mean of count per million (CPM) reads^57^) for co-occupancy of the H3K4me3 signal. Plots are centred on the TSS ± 3kb.

### ZBTB48 promotes open chromatin at CIITA pIII

Constitutive binding of ZBTB48 at CIITA pIII, even when the promoter is inactive (**Figure 1A**), suggests that ZBTB48-dependent promoter regulation primes transcriptional activation. To confirm that ZBTB48 binds independently of IFNγ, we performed ChIP-qPCR across all the CIITA promoters comparing each four U2OS WT and ZBTB48 KO clones both with and without IFNγ-treatment. In contrast to no detectable enrichment at the upstream CIITA pI, ZBTB48 binds strongly at CIITA pIII in the WT clones with >450-fold enrichment relative to the ZBTB48 KO clones (**Figure 5A**). In agreement with the notion of constitutive binding, this interaction was not altered upon induction of the promoter by IFNγ. Similar results were obtained for the MTFP1 promoter, a positive control that was established previously as a ZBTB48 regulated promotor irrespective of IFNγ^28^. Interestingly, we detected an additional relatively more moderate 17-fold enrichment of ZBTB48 compared to the ZBTB48 KO clones at the elusive CIITA pII, about 6-7kb upstream of pIII. Likewise, an 8-fold enrichment was also detected at pIV (**Figure 5A**). These comparably weak interactions might represent remnants of chromatin looping at the CIITA promoter region with CIITA pII acting as an enhancer element. Again, ZBTB48 binding at various loci within the CIITA locus is not affected by IFNγ, further validating that ZBTB48 itself does not respond to IFNγ.

Transcriptional programs primed for rapid induction are marked by pre-existing activating histone marks and accessible chromatin at promoters and enhancer regions. To ascertain ZBTB48-dependent reshaping of the promoter landscape, we compared histone marks at the CIITA promoter region in four U2OS WT and ZBTB48 KO clones each by ChIP-qPCR. Deposition of the H3K4me3 mark results in promoter activation^34^ and indeed, IFNγ treatment induced H3K4me3 marks at both CIITA pIII and pIV in U2OS WT but not ZBTB48 KO clones (**Figure 5B**). Again, using MTFP1 as a constitutively expressed gene regulated by ZBTB48, H3K4me3 marks were enriched at the MTFP1 promoter in the WT clones independent of IFNγ treatment (**Figure 5B**). We obtained similar results in SUDHL4 cells, a diffuse large B-cell lymphoma (DLBCL) cell line, where CIITA pIII is constitutively expressed. Again, loss of ZBTB48 led to a concomitant loss of H3K4me3 signal at both CIITA pIII and pIV (**Supplemental Figure S6A and 6B**). To test whether H3K4me3 is altered at promoters of other ZBTB48 target genes, we performed H3K4me3 ChIP-seq in three U2OS WT and ZBTB48 KO clones treated with IFNγ. Indeed, in agreement with the qPCR results, enrichment of H3K4me4 over CIITA pIII was observed in the WT clones (**Figure 5C, Table S3**). In addition, the H3K4me3 enrichment in WT cells extends to pIV, indicative of an active promoter status across the entire region. Concomitantly, downstream of CIITA induction, the majority of genes in the HLA cluster displayed robust H3K4me3 signal in U2OS WT cells but not in ZBTB48 KO clones (**Supplemental Figure S6D, Table S4**). However, not all promoters regulated by ZBTB48 depicted a similar loss of H3K4me3 levels in ZBTB48 KO cells despite a loss of target gene expression. For instance, the ZBTB48 target gene SNX15 retains its H3K4me3 signature at the proximal promoter in ZBTB48 KO cells despite a loss of gene expression (**Supplemental Figure S6C**). When globally aligning the ZBTB48 and H3K4me3 ChIP-seq data, the majority of ZBTB48 binding sites show strong concordance with H3K4me3 signal and the overall profiles remain constant between U2OS WT and ZBTB48 KO clones (**Figure 5D, Table S4**). Since only a subset of ZBTB48 binding sites respond to ZBTB48 deletion with a H3K4me3 reduction, ZBTB48 likely does not influence this histone modification directly. This is in agreement with the absence of H3K4me3 at CIITA pIII in uninduced U2OS WT cells despite constitutive binding of ZBTB48 to the promoter.

**Figure 6:**
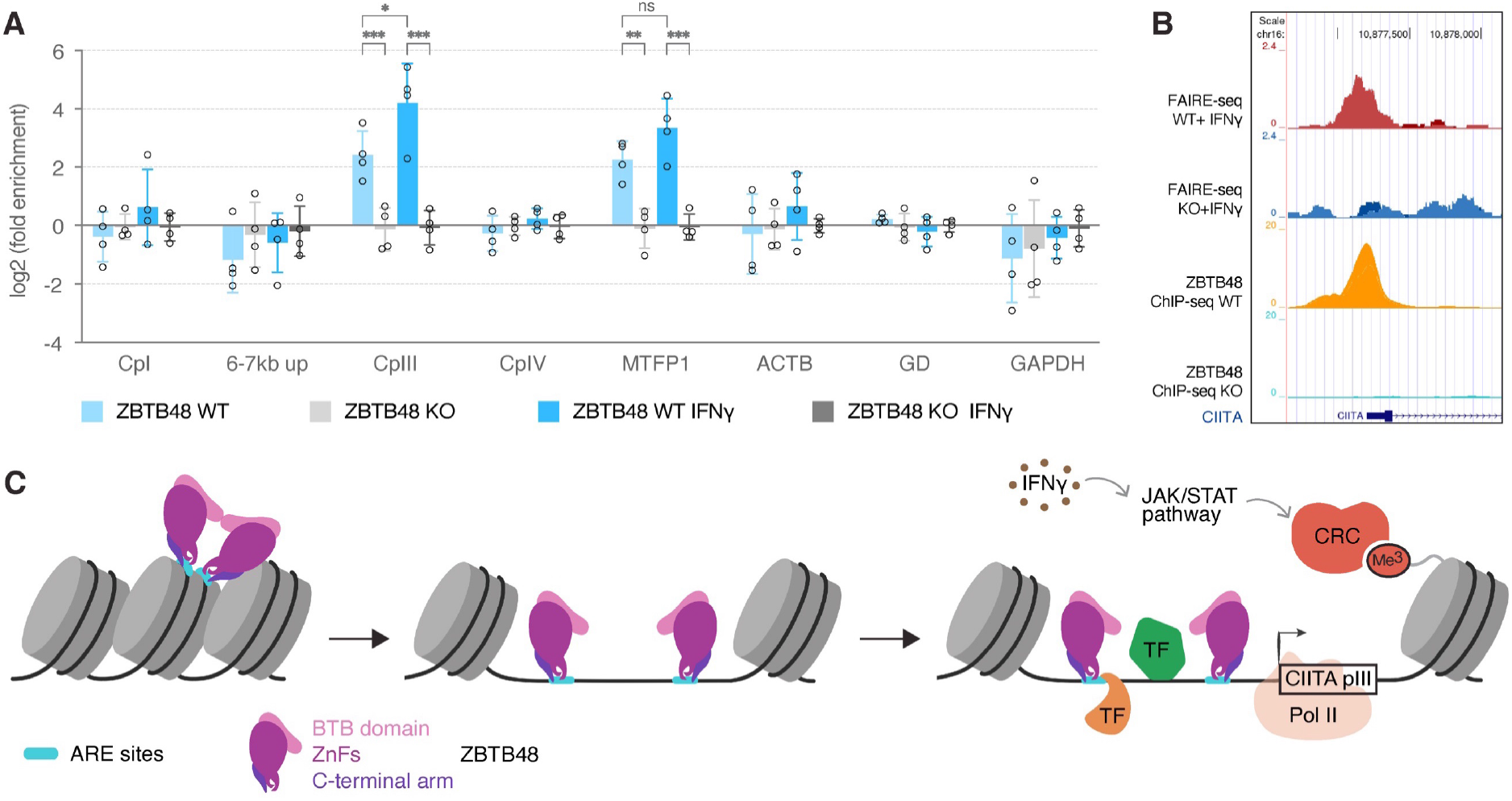
ZBTB48 establishes open chromatin at CIITA pIII. (A) FAIRE reactions from four independent U2OS WT and ZBTB48 KO clones with and without IFNγ induction analysed by qPCR for CIITA pI, pII (6-7kb up), pIII, pIV. MTFP1, GAPDH and ACTB (beta actin) promoters are used as a positive controls. Gene desert (GD) and a previously validated chromosome 12 region were used as negative controls. The data was normalised to the chromosome 12 region and fold-change was calculated relative to the average of WT clones. Data represents mean ± SD. p-values were calculated by 2-way ANOVA with Sidak correction for multiple comparisons (n = 4). (B) FAIRE-seq tracks depicting open chromatin peaks at CIITA pIII in two independent IFNγ-induced U2OS WT cells but not in the KO cells. The ZBTB48 ChIP-seq tracks are the same as described in Figure 1A. (C) Schematic illustration for the proposed role of ZBTB48 as a pioneering factor at CIITA pIII. The model depicts establishment of a nucleosome-depleted region at CIITA pIII upon binding of ZBTB48 binding at the ARE sites providing access to other transcription factors (TF) and chromatin remodelling complexes (CRC) upon IFNγ stimulation for transcription by pol II. Note that the two ARE binding sites would roughly be placed side-by-side in a nucleosome-bound state. The model was created with BioRender.

The above results suggest that ZBTB48 might act on chromatin regulation upstream of histone marks and hence we investigated chromatin accessibility. To test this at CIITA pIII, we performed Formaldehyde-assisted isolation of regulatory elements (FAIRE)^35^ followed by qPCR with the same promoter-specific primers used above in four WT and ZBTB48 KO clones as biological replicates. Chromatin accessibility at CIITA pIII, was 6-fold higher in the WT clones and this difference increased to 24-fold upon IFNγ treatment (**Figure 6A**). In comparison, the constitutively active MTFP1 promoter also displayed open chromatin in U2OS WT but not ZBTB48 KO clones and as expected this did not significantly alter upon IFNγ stimulation. CIITA pI, pIV and the 6-7kb upstream site were unaffected by the loss of ZBTB48, demonstrating that ZBTB48 specifically determines the chromatin state precisely at its binding site within CIITA pIII. To further establish this pioneering role of ZBTB48, we performed genome-wide FAIRE-seq analysis of open chromatin after IFNγ induction in U2OS WT and ZBTB48 KO clones. We examined the FAIRE peak positions with respect to the ZBTB48 binding locus at CIITA pIII, and indeed ZBTB48 binding peaks coincided with the open chromatin loci activation (**Figure 6B, Table S5**). Similar results were observed at other ZBTB48-dependent genes such as MTFP1 and SNX15 (**Supplemental Figure S6E and 6F**), suggesting that ZBTB48 in general ensures chromatin accessibility at the promoters of its targets.

Based on the above, we therefore propose that ZBTB48 functions as a pioneer factor that sets the stage for promoter activation (**Figure 6C**). In such a model, the binding of ZBTB48 at nucleosome-compacted promoters induces a nucleosome-depleted region to subsequently grant access to other transcription factors and chromatin remodelling complexes. In the case of CIITA, this requires additional IFNγ stimulation and binding by IFNγ-dependent transcription factors, which in turn solidify the open chromatin state and trigger transcription initiation by RNA polymerase II.

## DISCUSSION

The precisely fine-tuned and strictly regulated cell type specific expression of MHC II is primarily regulated at the transcriptional level by the differential usage of the three independent promoters of CIITA. In addition to being IFNγ-inducible, pIII predominantly drives the constitutive expression of CIITA in B cells. Here, we describe ZBTB48 as a key regulator of both IFNγ-inducible and constitutive *in vivo* expression of CIITA pIII. Prior work using gene reporter assays had established that ARE-1 and ARE-2 are the essential regulatory elements for CIITA pIII^22, 25^. Intriguingly, for at least ARE-2, the authors had also implicated the GGG motif as critical in refined promoter mutants^25^. Our serendipitous discovery of ZBTB48 hence ideally fits in this picture with ZBTB48’s ability to recognise GGG motifs via ZnF11. While other ARE-binding proteins, including AML2/3, CREB1 and ATF1, had been previously suggested^22, 25–27^, these data are limited to supershifts in EMSA assays and therefore did not include *in vivo* validation. It remains to be evaluated whether any of these proteins specifically compete with ZBTB48 for binding to the ARE elements. Nonetheless, the near complete absence of CIITA pIII induction in U2OS ZBTB48 KO clones suggests that at least in this cellular context ZBTB48 is the essential ARE transducer.

While we observed similarly striking phenotypes in pre-B cells in our ZBTB48 KO mouse model, the MHC II loss was less pronounced in mature B cells as well as in earlier and later B cell developmental stages. These data suggest that there is heterogeneity in the B cell population either in their CIITA promoter choice and/or in terms of expression of potentially redundant regulators. Even in systemic CIITA knock- out mice, 1-3% MHC II-positive splenic B cells were detected^36^, implying that the small portion of ZBTB48-independent, MHC II-positive pre-B cells might be selected for and populate the mature B-cell stages. While at present we cannot explain why only specifically female ZBTB48^-/-^ mice present with a mild splenomegaly, quality control mechanisms triaging out MHC II-negative precursors, might contribute to the observed phenotype. It will be interesting to see whether at later time points splenomegaly starts appearing in males as well and whether it overall aggravates with age. In addition, one would predict that such a scenario should ultimately lead to stem cell exhaustion in the B-cell lineage with increasingly smaller numbers of MHC-positive cells in mature B cell populations and/or a reduction in the corresponding cell numbers. However, in contrast to the above, systemic CIITA knock-outs overall lack cell surface MHC II expression on splenic B cells and DCs, and their macrophages and somatic tissue cells also do not express MHC II upon IFNγ induction^36, 37^. Likewise, mice specifically deficient in p(III+IV) lack MHC II expression in all B cell subsets, including in the spleen and blood^38^. Since studies in mice targeting exclusively pIII are lacking, a direct comparison with our results is impossible at present, but these data imply that exclusive loss of pIII-driven CIITA can be partially compensated at least in young animals.

Molecularly, our results suggest that ZBTB48 acts as a pioneer factor that is essential to activate the proximal promoters of its target genes. Intriguingly, the two ZBTB48 binding sites at their GGG motif core are ∼80 nucleotides apart from each other (**Figure 1D**), which in a nucleosome context places both binding sites more or less exactly in adjacent positions^39^. Given that many ZBTB family members are known to homo- or hetero-dimerize via their BTB domain^40^, it is tempting to speculate that recognition of nucleosome bound CIITA pIII is achieved by a ZBTB48 dimer. Upon opening of the locus, CIITA pIII would subsequently become accessible for other transcription factors that in turn are required to trigger a cascade of gene expression (**Figure 6C**). While the inducible nature of CIITA pIII allows us to distinguish such a step-wise process, we envision a similar control at other ZBTB48 target genes with the companion transcription factors being constitutively expressed. In such a view, ZBTB48 could be equated to an on-off light switch while additional transcription factors function as dimmer switches. Hence, while ZBTB48 determines whether gene activation is possible at all, the exact transcriptional output is regulated downstream of the pioneering activity by other transcription factors.

In a similar vein, it will be interesting to see how much of these mechanistic details translate from ZBTB48’s transcription factor function to its role at telomeres. Prior work had established that ZBTB48 associates more frequently with telomeres of cancer cells relying on the Alternative Lengthening of Telomeres (ALT) mechanism^28, 29, 41^. This telomere maintenance mechanism is in parts initiated by epigenetic changes downstream of loss of ATRX or DAXX^42–45^ and depletion of these chromatin regulators in normal cells paves the way for an increased association of ZBTB48 with telomeres^41^. Beyond potentially providing a permissive chromatin environment for ZBTB48-telomere association, it will be important to determine whether ZBTB48 itself regulates telomere compaction. Likewise, these dual functions provide an exciting additional connection between telomere biology and immune regulation and it will be interesting to clarify the crosstalk between both processes. At the same time, the nuanced difference in ZBTB48’s binding behaviour, with ZnF10 being essential for CIITA but not telomeric recognition, provides a clearly defined separation-of-function mutant. This will allow future research to carefully distinguish in cis from in trans effects.

Finally, ZBTB48 differs from previously studied pioneering factors, for which the prevailing expectation is lineage-specific expression that supervises cell-specific epigenomic profiles^46, 47^. In contrast, ZBTB48 expression itself does not show predominant tissue-specificity. In addition, while the effect of ZBTB48 on its target genes is almost all or none, the number of differentially expressed genes between ZBTB48 WT and KO clones is relatively small. Given that the canonical C2H2 zinc finger-containing genes alone already sum to more than 800 family members in the human genome^48^ and only a small portion of protein-coding genes make up the majority of the scientific literature, it seems conceivable that many zinc finger proteins may follow ZBTB48’s pattern. Having a large number of transcription factors that function as on-off-switches for a handful of target genes would provide exquisite specificity upstream of dimmer switch regulation that impacts the relative transcriptional output. Such a model would likely include redundancy and hence genes that can be activated by multiple transcription factors. This in return would explain the larger number of proximal promoter binding sites at genes that are not affected in their gene expression profile in ZBTB48 KO cells. At such a systems biology level, the coordinated effort of on-off-switch regulators would thus provide an underlying pattern to orchestrate a logic expression circuit.

## METHODS

### Cell culture

U2OS and HEK293T were cultured in Dulbecco’s Modified Eagle’s Medium (DMEM) containing 4.5 g/l glucose, 4 mM glutamine, 1 mM sodium pyruvate and supplemented with 10% foetal bovine serum (FBS; Gibco), 100 U/ml penicillin and 100 μg/ml streptomycin (Gibco). SUDHL4 DLBCL cells were cultivated in RPMI-1640 (Gibco) medium supplemented with 10% FBS, 2 mM glutamine, 100 U/ml penicillin and 100 μg/ml streptomycin. All three cell lines were maintained in a humidified incubator at 37°C and 5% CO_2_. For induction of CIITA, 250 U/ml IFNγ was added to the culture medium and cells were incubated for 24 and 48 h for RNA and protein extraction, respectively.

### Deletion mutant/variant construction

ZBTB48 zinc finger (ZnF) point mutants and deletion variants had been established previously^28^. pcDNA3.1-FLAG-ZBTB48 WT was used as template for the generation of the constructs with point mutations in the C-terminal arm using the primers in **Table S6** and the Q5 Site-Directed Mutagenesis Kit (NEB). In brief, exponential amplification was performed with Q5 Hot Start High-Fidelity Master Mix according to the manufacturer’s protocol using 10 ng of template with the following cycling conditions on a SimpliAmp thermal cycler (Applied Biosystems): 98°C for 30 s, followed by 25 cycles with 98°C for 10 s, 67°C for 30 s, 72°C for 3 min and a final extension at 72°C for 3 min. The PCR product was phosphorylated and ligated with KLD enzyme mix (NEB) before transformation into competent cells followed by sequence verification (1st BASE) of the constructs.

### Transfection

Plasmids were transfected in U2OS cells using linear polyethylenimine (PEI, MW 25,000; Polysciences). 15.2 μg of plasmid and 60.8 μg of PEI were diluted in 2ml of Opti-MEM, mixed and incubated at room temperature for 20 min. The transfection mix was incubated with 4.5 × 10^6^ cells seeded in a 15 cm dish for 7 hours before washing with 1x PBS and replacement with fresh culture medium. The cells were collected 48 hours post transfection for nuclear protein extraction.

For lentiviral packaging in HEK293T cells, each lentiviral transfer vector (0.5 μg) was co-transfected with 0.25 μg each of pMDLg/pRRE and pRSV-Rev and envelope vector pMD2.G (0.25 μg) into 3 × 10^5^ cells in a 6-well plate. The plasmids were combined with 2.5 μg PEI in 100 μl Opti-MEM and incubated at room temperature for 20 min before incubating with cells for 18 hours. The cells were washed with 1x PBS and replenished with fresh culture medium. Lentiviral supernatants were collected 48 hours after transfection and filtered with 0.22 μm syringe filters and stored at -80°C until use for transduction.

### Nuclear protein extraction

Nuclear extracts were prepared by incubating cells in hypotonic buffer (10 mM Hepes, pH 7.9, 1.5 mM MgCl_2_, 10 mM KCl) on ice for 10 min. Cells were transferred to a dounce homogenizer in hypotonic buffer supplemented with 0.1% Igepal CA630 (Sigma) and 0.5 mM DTT and lysed by 40 strokes. Nuclei were washed once in 1x PBS and incubated in hypertonic buffer (420 mM NaCl, 20 mM Hepes, pH 7.9, 20% glycerol, 2 mM MgCl_2_, 0.2 mM EDTA, 0.1% Igepal CA630 (Sigma), 0.5 mM DTT) for 2 hours at 4 °C on a rotating wheel. Nuclear extract was collected by centrifugation at maximum speed for 1 hour at 4 °C. Protein amounts were quantified with the Pierce BCA Protein Assay Kit according to the manufacturer’s instructions (Thermo Scientific).

### DNA pulldown

Equal amounts (25μg) of forward and reverse sequence oligos (**Table S6**) corresponding to each of the probes were combined in annealing buffer (20 mM Tris– HCl, pH 7.5, 10 mM MgCl_2_, 100 mM KCl), denatured at 80°C, and annealed by cooling. 100 units T4 kinase (Life Technologies) was added and incubated for 2 hours at 37°C for phosphorylation, followed by overnight incubation with 20 units T4 ligase. The concatenated DNA strands were purified using phenol–chloroform extraction, biotinylated with desthiobiotin-dATP (Jena Bioscience) and 60 units DNA polymerase (Thermo Scientific) and finally purified using microspin G-50 columns (GE Healthcare). The probes generated were immobilized on 250 μg paramagnetic streptavidin beads (Dynabeads MyOne C1, Thermo Scientific) on a rotation wheel for 30 min at room temperature. Subsequently, baits were incubated with 75 μg nuclear extract in PBB buffer (150 mM NaCl, 50 mM Tris–HCl pH 7.5, 5 mM MgCl_2_, 0.5% Igepal CA-630 (Sigma) and 1 mM DTT) while rotating for 2 hours at 4°C. As a competitor for DNA binding, 20 μg sheared salmon sperm DNA (Ambion) was added. After three washes with PBB buffer, bound proteins were eluted in 2× Laemmli buffer (Sigma-Aldrich), boiled for 5 min at 95°C and separated on a 4-12% NuPAGE Novex Bis-Tris precast gel (Thermo Scientific).

### Quantitative RT-PCR

RNA was isolated using the RNeasy Plus Mini Kit (Qiagen) and 1 μg of RNA was used for cDNA synthesis using the SuperScript IV first-strand synthesis kit with oligo-dT following the manufacturer’s protocol. Quantitative PCR (qPCR) was carried out using 1x QuantiNova SYBR green PCR mix (Qiagen) and 500 nM each of the forward primer and reverse primer (Table 3) with the following cycling conditions on either QuantStudio 3 or 5 Real-Time PCR systems (Applied Biosystems): 95°C for 2 min, followed by 40 cycles with 95°C for 5 s, 60°C for 30 s followed by a melting curve. TBP was used as a housekeeping gene for normalization. All data were analysed using Prism 8 (GraphPad Prism, La Jolla, CA).

### Western blot

Protein samples were denatured in 2x Laemmli buffer (Sigma-Aldrich) at 95°C for 5 min and separated on a 4-12% Bis-Tris gel (NuPAGE, Thermo Scientific) for 60 min at 170V. For detection of CIITA, cells were lysed with 1x RIPA buffer supplemented with 1x cOmplete proteinase inhibitor (Roche) for 30 min on ice, pelleted and denatured with 1x LDS sample buffer (Thermo Scientific) at 95°C for 5 min in the presence of 100 mM DTT. Protein amounts were quantified with the Pierce BCA Protein Assay Kit (Thermo Scientific) and at least 50 μg of protein were used. Samples were transferred to a PVDF membrane (Bio-Rad) using a semi-dry transfer system for 90 min at 50 mA per blot with 1x Transfer Buffer (25 mM Tris, 192 mM glycine, 0.02 % (v/v) SDS, 10 % (v/v) methanol). The membrane was blocked in PBS containing 0.1% Tween-20 (PBST) and 5% (w/v) non-fat milk for 1 hour at RT prior to incubation with primary antibody (**Table S7**) overnight at 4°C. After three washes in 5% milk PBST for 10 min each at RT, the membrane was incubated for 60 min at RT with secondary antibodies (**Table S7**) in 5% milk PBST, followed by two washes in 5% milk PBST, one wash in PBST 10 min each at RT and finally rinsed with 1x PBS before detection. The membrane was revealed with ECL substrate (Pierce ECL Western blotting substrate (Thermo Scientific) or Amersham ECL Prime Western blotting detection reagent (GE Healthcare) using either a ChemiDoc imaging system (Bio-Rad) or X-ray films. PageRuler Plus Prestained Protein Ladder (Thermo Scientific) and MagicMark XP Western Protein Standard (Thermo Scientific) were used as size markers.

### Lentiviral transduction for generation of knockout cells

U2OS ZBTB48 WT and KO clones used here have been previously described^28^. In SUDHL4, CRISPR/Cas9 mediated ZBTB48 deletions were performed using three different sgRNA sequences (**Table S6**) targeting its second exon. The sgRNAs were cloned into lentiCRISPRv2-puro vector and confirmed by Sanger sequencing (1st BASE). The lentiCRISPRv2-puro vector containing the Cas9 expression cassette and the gRNA scaffold was a gift from Brett Stringer (Addgene plasmid #98290 ; http://n2t.net/addgene:98290 ; RRID:Addgene_98290). For transduction, 700 ul of lentiviral supernatant, generated using HEK293T cells and supplemented with 10 mM HEPES and 8 µg/mL polybrene (Sigma Aldrich), was combined with 100 μl of cell suspension containing 5*10^5^ SUDHL4 cells in 12 well plates and spin-inoculated by centrifugation at 2000 rpm for 1.5 hours at 32°C. The reaction mix was replaced 24 hours post transduction with fresh culture medium. Selection with 500 ng/ml puromycin was performed 48 hours after transduction, over 2 weeks before T7 endonuclease assay (T7E1) assay. Upon confirmation of successful cleavage by Cas9, cells were single sorted using a BD FACS Aria II flow cytometer and expanded to screen for ZBTB48 KO clones by Western blot. Similarly, SUDHL4 WT cells were also single sorted and used as WT clones. The genotype of clones with undetectable levels of ZBTB48 and its downstream target, MTFP1, was further confirmed by Sanger sequencing. PCR products, amplified using gDNA of each clone using the same amplification protocol as the T7E1 assay, were cloned into the pCR2.1-TOPO TA vector (TOPO TA cloning kit, Thermo scientific) following the manufacturer’s instructions and ten colonies from each clone were sequenced (1st BASE).

### T7E1 assay

A T7 endonuclease I assay was performed to evaluate Cas9 induced mutations. Genomic DNA was extracted from puromycin selected SUDHL4 cells and untreated WT cells using the QIAamp DNA Blood Mini Kit (Qiagen) following the manufacturer’s instructions. Using 100 ng of gDNA, a 835bp region around the cut locus was amplified using the Q5 Hot Start High-Fidelity DNA Polymerase (NEB) and 400nM of each primer (**Table S6**) with the following PCR conditions: 94°C for 3 min followed by 35 cycles with 94°C for 30 s, 60°C for 30 s, 72°C for 50 s and a final extension at 72°C for 5 min. PCR products were analysed by gel electrophoresis and purified using QIAquick PCR purification kit (Qiagen). 100 ng purified DNA each from WT and sgRNA treated samples were denatured at 95°C for 5 min in 1× NEB 2 buffer and hybridized following the temperature profile 95 to 85°C at 2°C/s and 85 to 25°C at 0.1°C/s. For digestion of heteroduplexes, 2 μl of 10× NEB 2 buffer, 1 μl T7 Endonuclease I (NEB) and 7 μl HPLC-grade H2O were added, and samples were incubated at 37°C for 20 min. Cas9 activity was confirmed by separation of products on a 2% agarose gel.

### Generation of knockout mice

C57BL/6NTac and CD1 mice were obtained from Invivos, Singapore and Charles River Laboratories, USA respectively. All animal protocols were approved by the Institutional Animal Care and Use Committee (IUCAC) at the National University of Singapore, Singapore (protocol number: BR20-0926 and R20-0927). Animals were housed with food and water ad libitum, on a 12 hour light/dark cycle (lights off at 7pm) and the health status of the animals was routinely checked by a veterinarian.

Two sgRNAs (**Table S6**) targeting regions at the opposite ends of exon 2 were designed using CRISPick (Broad Institute). The sgRNAs (Integrated DNA technologies, Singapore (IDT)) were tested in NIH3T3 mouse cell line prior to their use in the mice. CRISPR/Cas9 mediated transgenic mice were generated with the help from the Transgenic and Gene Targeting Facility at CSI, NUS. Briefly, equimolar amounts of crRNA and tracrRNA were annealed in microinjection buffer (MIB; 1mM Tris, 0.25 mM EDTA ph 7.4) following the IDT protocol. The RNP (ribonucleoprotein) complex was prepared by incubating the reaction mix with equimolar amount of Alt-R S.p. Cas9 nuclease (# 1074181, IDT, Singapore) for 5 minutes at room temperature in MIB and then combined with ssODN to prepare the microinjection cocktail. Single cell staged C57BL/6NTac mouse embryos were harvested from superovulated 4 week old female mice at 0.5 d postcoitum after mating with male mice of the same strain^49^. The microinjection cocktail was injected into either one of the pronuclei of C57BL/6NTac mouse embryos with Leica AM6000 manipulator using standard protocol^49^. The zygotes were cultured overnight in KSOM mouse embryo culture medium and twenty five viable zygotes were transferred to oviducts of 0.5 d postcoitum psedopregnant female CD1 mice. The pup tail clippings were genotyped by PCR following the protocol below. All transgenic mouse lines were back-crossed into C57BL/6NTac background and experiments were performed on 8-14 week old mice using littermate trios of -/-, +/-, +/+ of same sex.

### Mouse genotyping

2-3 week old mice were ear tagged and tail clippings were collected for genotyping. For gDNA isolation, each tail tip was incubated in 500 μl of Lysis buffer (200m M NaCl, 100 mM Tris–HCl pH 8.5, 5 mM EDTA, 0.2% SDS) with 2.5 μl of Proteinase K (20mg/ml) overnight at 55°C under constant agitation (800 rpm). 500 μl of digestion solution was added to the digestion solution and the precipitated DNA was fished out with a pipette tip and dissolved in 1x TE buffer for 2 hours at 55°C. Using 400nM each of primers that flank the cut sites (**Table S6**) PCR was performed using 1 μl of dissolved DNA, 1x MyTaq Red Reaction buffer red and 0.5 μl of MyTaq DNA polymerase (Bioline) on SimpliAmp thermal cycler (Applied Biosystems) using the following cycling conditions: 94°C for 3 min followed by 35 cycles with 94°C for 30 s, 60°C for 30 s, 72°C for 50 s and a final extension at 72°C for 5 min. The genotype was determined based on the product size of the product by separation on a 1% agarose gel for 60 min at 100 V.

### Isolation and sorting of mouse cells

Retro-orbital bleeding was performed to collect 20 µl of blood from 8-12 week old mice. The mice were euthanised, and spleen, thymus, long bones (tibias and femurs) and spine were collected for analysis. Viable single cell suspension from spleen and thymus was obtained by rubbing small pieces of the organ through a nylon filter (40 µm). BM cells were flushed from the long bones and spine with 1x PBS and next filtered through a nylon filter (35 µm) to obtain a single-cell suspension. The red blood cells from the cell suspension were lysed with Ammonium Chloride Potassium (ACK) red cell lysis buffer (150mM Ammonium chloride, 10mM Potassium bicarbonate, 0.1 mM EDTA) for 5 min at 4°C. The lysis reaction was stopped by diluting the cell suspension mix to ten times the volume with 1x PBS and the cells were pellet by centrifugation at 1000 rpm for 5min and subsequently resuspended in 1x PBS for sorting and analysis. The RBC lysis step was performed twice for blood samples.

### Fluorescence activated cell sorting (FACS) analysis

Hematopoietic stem and progenitor cells, and lineage positive cells from BM, spleen and thymus of mice from each genotype were profiled by FACS. Littermates with at least one each of WT, Het and KO were analysed together. Prior to antibody staining, the cells were blocked with FcgR (CD16/32) for 15 min on ice and then incubated with antibodies (**Table S7**) for additional 20 min on ice. 0.25 ng/mL of Hoechst 33258 was used to exclude dead cells. Samples were analyzed on a BD LSRII Flow Cytometer or a BD FACSAria cell sorter.

### Protein expression and Crystallization

The ZBTB48 ZnF10-11-C construct containing ZnF10-11 and the conserved C-terminal region (residues 548-620) was amplified from human brain cDNA library and cloned into a modified pGEX-4T-1 vector containing TEV (Tobacco Etch Virus) cleavage site. The proteins were induced in Escherichia coli BL21 (Gold) cells with 0.1 mM isopropyl β-D-1-thiogalactopyranoside (IPTG) for 5 hour and purified by glutathione sepharose (GE healthcare) followed by on-column TEV cleavage and finally by size-exclusion chromatography on Hiload 16/60 Superdex 75 coloumn (GE healthcare).

The purified ZnF10-11-C protein was combined with double-stranded CIITA e.2 probe sequence (G-strand: 5’-CACAAGTGAGGGATCA-3’; C-strand: 5’- GTGATCCCTCACTTGT-3’) at a 1:1.5 molar ratio in buffer containing 20 mM Tris-HCl (ph 7.5) and 500 mM NaCl. The reaction mix was dialyzed against a buffer containing 20 mM Tris-HCl (pH 7.5) and 150 mM NaCl and concentrated to 1 mM. Using the hanging drop vapor diffusion method, the crystals were grown at 20°C by combining equal volumes of the protein-DNA complex and reservoir buffer (0.2 M imidazole malate, pH 8.5 and 20 % (w/ v) polyethylene glycol 4K). Reservoir buffer with 25% glycerol was used as cryoprotectant.

X-ray diffraction data collection was performed on beamline 19U1 (BL19U1) at the Shanghai Synchrotron Radiation Facility (SSRF) at wavelength 0.9758 and *HKL2000* software (HKL Research)^50^. The initial crystallographic phases were calculated using molecular replacement which was carried out by Phaser employing the ZBTB48- telomeric DNA complex structure as the search model (PDB code: 5YJ3)^51, 52^ and the model was further built and refined using Coot and Phenix.refine^51, 53^.

### Next-generation RNA sequencing and data analysis

RNA was extracted from IFNγ (250U/ml, 24 hours) treated and untreated control U2OS WT and ZBTB48 KO clones using the RNeasy Plus Mini Kit (Qiagen) with additional on-column DNaseI digestion following the manufacturer’s instructions. Five clones per condition were used as biological replicates. RNA quality assessment, library preparation and sequencing was performed at NovogeneAIT (Singapore). RNAseq (stranded mRNA) library prepared after quality assessment (RIN#>7) were sequenced on a NovaSeq 6000 PE150 and 53 to 82 million reads were obtained per sample. Reads were aligned to the human reference genome version GRCh38 (GCA_000001405.15) using STAR v2.7.9a^54^ whose index was generated with RefSeq annotation and up to 4 mismatches were allowed per pair. The alignments were filtered for unique mappers using NH:i:Nmap field with value 1 and further quantified using featureCounts v2.4.0^55^. DESeq2 was used for differential expression analysis and log fold changes were shrunk using the original DESeq2 shrinkage estimator (normal)^56^. Bigwig tracks were generated using bamCoverage (deeptools v3.5.1)^57^, hosted on cyverse^58^ and visualized on UCSC Genome Browser^59^.

### MS sample preparation

Protein samples were boiled in Laemmli buffer (Sigma-Aldrich) at 95°C for 5 min and separated on a 12% Bis-Tris gel (NuPAGE, Thermo Scientific) for 30 min at 170 V. Proteins were stained using the Colloidal Blue Staining Kit (Thermo Scientific) according to the manufacturer’s instructions. For in-gel digestion, samples were separated in 4 equal gel pieces for each sample from high to low molecular weight and each fraction was cut individually with a clean scalpel into 1 mm x 1 mm pieces. First, gel pieces were destained in destaining buffer (50% 50 mM ammonium bicarbonate buffer, 50% ethanol) twice followed by two rounds of dehydration in 100% acetonitrile. Supernatants were discarded and residual liquid removed using a Concentrator Plus (Eppendorf). Samples were then reduced in 10 mM DTT (Sigma) for 1 h at 56°C followed by alkylation with 55 mM iodoacetamide (Sigma) for 45 min in the dark. Tryptic digest was performed in 50 mM ammonium bicarbonate buffer with 2 μg sequencing-grade trypsin (Promega) at 37°C overnight. Supernatants were collected and the digested peptides extracted from the gel pieces by one round of extraction buffer (30% acetonitrile, 10% trifluoracetic acid, 70% mM ammonium bicarbonate buffer), one round of 100% acetonitrile, another round of extraction and two more rounds of 100% acetonitrile for 15-20 min at RT with shaking. All supernatants were combined and evaporated in a Concentrator Plus (Eppendorf) to reduce the volume to < 200 μl. Peptides were desalted on self-made C18 Stage Tips and analysed with a Q Exactive HF mass spectrometer (Thermo) coupled to an EASY-nLC 1200 system (Thermo) nanoflow liquid chromatography system. Peptides were separated on a C18 reversed-phase capillary (25 cm long, 75 μm inner diameter, packed in-house with ReproSil-Pur C18-AQ 1.9 μm resin (Dr. Maisch) directly mounted on the electrospray ion source. We used a 215-min gradient from 2 to 40% acetonitrile in 0.5% formic acid at a flow of 225 nl/min. The Q Exactive HF was operated in positive ion mode with a Top20 MS/MS spectra acquisition method per MS full scan. MS scans were obtained with 60,000 resolution at a maximum injection time of 20 ms and MS/MS scans at 15,000 resolution with maximum IT of 75 ms.

### MS data acquisition and data analysis

The raw files were processed with MaxQuant^60^ version 1.5.2.8 against the UNIPROT annotated human protein database (81,194 entries). Carbamidomethylation on cysteines was set as fixed modification while methionine oxidation and protein N-acetylation were considered as variable modifications. Peptide and protein FDR were enforced at 0.01 with match between runs option activated. MaxLFQ quantitation^61^ was based on unique peptides for protein groups with at least 2 ratio counts. The resulting protein groups table was further processed with an in-house script to exclude contaminants (based on the MaxQuant contaminant list), reverse hits and proteins that were only identified by a modified peptide. Missing values were imputed based on normally distributed values corresponding to the percentage of missing values in each sample for proteins that were quantified in at least 4 out of 5 biological replicates.

### Chromatin immunoprecipitation (ChIP) and quantitative PCR

Cells were washed twice with ice cold 1x PBS and crosslinked for 20 min at RT with 1% (v/v) formaldehyde (Pierce) diluted in respective cell culture media. The reaction was quenched by addition of Glycine-PBS (0.125 M final concentration) for 5 min at RT and washed with ice cold 1x PBS. The fixed cells were collected by scraping in 0.5 ml of ice cold 1x PBS supplemented with 1x cOmplete proteinase inhibitor (Roche) for U2OS and by centrifugation for SUDHL4 at 1000g for 5 min at 4°C. The pellets were snap frozen and stored at in -80°C until lysis. For lysis, cells were resuspended and incubated at 4°C on a rotating wheel in Lysis buffer 1 (50 mM Tris–HCl pH 8.0, 250 mM sucrose, 140 mM NaCl, 1 mM EDTA pH 8.0, 10% glycerol, 0.5% Igepal CA-630, 0.25% Triton X-100, 0.25% Tween-20, 1x cOmplete proteinase inhibitor) for 15 min, followed by Lysis buffer 2 (10 mM Tris–HCl pH 8.0, 200 mM NaCl, 1 mM EDTA pH 8.0, 0.5 mM EGTA pH 8.0, 1x cOmplete proteinase inhibitor) for 10 min with centrifugation at 1000g for 10 min at 4°C in between. The nuclear pellet was collected by centrifugation and resuspended in sonication buffer (1% SDS, 10 mM EDTA pH 8.0, 50 mM Tris–HCl pH 8.0, 1x cOmplete proteinase inhibitor). The chromatin in sonication buffer was sheared to 200-400bp by sonication using an EpiShear Probe Sonicator (Active Motif) at 30% amplitude, 25 cycles of 15s ON/ 30s OFF for U2OS and 30 cycles for SUDHL4. Lysates containing 25 μg of sheared chromatin were incubated with 0.833 μg of antibodies (**Table S7**) in PBB1 buffer (180 mM NaCl, 50 mM Tris-HCl pH 7.5, 5 mM MgCl_2_, 1mM DTT, 0.5% IGEPAL CA-630, 1x cOmplete proteinase inhibitor) overnight at 4°C on a rotating wheel. Prior to use, for each ChIP reaction 12.5 μl of Dynabeads protein G magnetic beads (Thermo Scientific) were blocked with 33.3 μg of sheared salmon sperm DNA (Thermo Scientific) in PBB1 containing 10 μg/μl BSA for 10 min at room temperature on rotating wheel. The pre-blocked beads were mixed with the ChIP reactions containing antibodies and incubated for 2h at 4°C on a rotating wheel followed by seven washes with PBB2 buffer (180 mM NaCl, 50 mM Tris-HCl pH 7.5, 5 mM MgCl_2_, 1mM DTT, 0.5% IGEPAL CA-630, 1x cOmplete proteinase inhibitor) and once with ice cold 1x TE buffer. DNA was eluted twice in 100 μl of elution buffer (0.1 M NaHCO_3_, 1% SDS) and de-crosslinked overnight at 65°C in the presence of 0.2 M NaCl. The DNA samples were treated with 0.5 μg/μl of RNase A for 30 min at 37°C and then with 0.25 μg/μl of Proteinase K (Roche) in the presence of 10mM EDTA, 40mM Tris-HCl ph 6.5 for 2 hour at 45°C prior to purification with QIAquick PCR Purification Kit (Qiagen). For ‘input’ DNA, 2.5 μg (10% of ChIP reaction) of sheared chromatin lystes were treated similarly to elutes. qPCR was performed using ChIP primers indicated in Table 7 with the same amplification protocol mentioned in the qPCR section.

Next-generation ChIP sequencing (ChIP-seq) was performed by NovogeneAIT (Singapore). The purified DNA samples were assessed for quality prior to library preparation and sequenced on a NovaSeq 6000 PE150 and 41 to 53 millions of 150bp paired end reads were obtained per sample. Reads were trimmed for adapter sequences using trimmomatic PE v0.39^62^ and aligned to the human reference genome version GRCh38 using bowtie2 version 2.3.5.1^63^ with default settings and processed (sorting and indexing) using samtools version 1.12^64^. Multimapping and unmapped reads were filtered out and duplicate reads were removed using sambamba version 1.0^65^. Peaks were called using MACS version 2.1.1.2^66^ in paired-end mode with default q-value cut-off (0.05) and --broad flag. We performed a quantitative differential binding analysis with Diffbind 3.2.1^67^ between the WT and ZBTB48 KO conditions. Peaks which were significantly different (FDR < 0.01 and fold change > 2) and called consistently in all 3 WT replicates were marked as “TRUE” in the “DB” column. The resulting peak set was annotated with the closest TSS using the hg38 annotation database generated from UCSC. Bigwig tracks normalized to counts per million mapped reads were generated using bamCoverage (deeptools v3.5.0)^57^, hosted on cyverse^58^ and visualized on UCSC Genome Browser^59^.

### Formaldehyde-assisted isolation of regulatory elements (FAIRE)

FAIRE was performed by phenol-chloroform extraction of the sonicated 1% formaldehyde fixed samples obtained as described in the ChIP section. For collection of ‘free chromatin’ 30 μg of sonicated samples was treated with 10 μg/μl of RNase A (Thermo Scientific) in the presence of 0.2 M NaCl at 37°C for 1.5 hours. DNA was obtained after two extractions with phenol-chloroform-isoamyl alcohol followed by a wash with chloroform followed by precipitation in the presence of 0.5 M NaCl, 40 μg of glycogen and equal volume of isopropanol at -80°C for at least 1 hour. After final precipitation by 70% ethanol, the DNA pellet was resuspended in 30 μl of nuclease-free water and incubated at 65°C overnight for de-crosslinking. Input DNA was obtained following the sample phenol-chloroform purification steps post overnight de- crosslinking of 10 μg of sonicated sample at 65 °C in the presence of 0.2 M NaCl and 10 μg/μl of RNase A and 0.8 μg/μl of Proteinase K (Roche). DNA was quantified with the Qubit dsDNA HS Assay Kit (Thermo Scientific) according to manufacturer’s instructions. qPCR was performed with 10ng of DNA using the same primers and amplification protocol as for ChIP (**Table S6**) and relative free DNA was estimated as outlined by Rodriguez-Gil *et al*^35^.

For FAIRE-seq, free chromatin and input DNA was assessed for quality prior to library preparation and sequencing at NovogeneAIT (Singapore). Sequencing was performed on a NovaSeq 6000 PE150 and 41 to 62 millions of 150bp paired end reads were obtained per sample. Reads were stripped of adapter sequences with trimmomatic PE v0.39^62^ and mapped to human reference genome (GRCh38) using bowtie2 v2.2.5^63^, while reads that had a mapping quality less than 30 were removed using samtools v1.14^64^. Peaks were called using MACS 2.2.7.1^66^ in paired-end mode and subject to irreproducible discovery rate analysis (IDR 2.0.4.2)^68^ wink rank based on p-value and allowing non-overlapping peaks to measure reproducibility between replicates. Further, reproducible regions (IDR score > 125) in replicates of U2OS WT and ZBTB48 KO respectively were considered to determine areas of enrichment by calculating the RPKM and log2 foldchange at these regions using read counts reported by multiBamCov (bedtools v2.30.0)^69^. Bigwig tracks normalized to counts per million mapped reads were generated using bamCoverage (deeptools v3.5.1), hosted on cyverse^58^ and visualized on UCSC Genome Browser^59^.

## ACKNOWLEDGMENTS

We are grateful to all members of the Kappei lab for advice and discussions. This research was supported by the National Research Foundation Singapore and the Singapore Ministry of Education under its Research Centres of Excellence initiative, funding by the Singapore Ministry of Education’s Tier 3 grants (MOE2014-T3-1-006), funding by the Fritz Thyssen Foundation (10.21.1.022MN) and an NMRC Open Fund Individual Research Grant (MOH-OFIRG21jun-011).

## AUTHOR CONTRIBUTIONS

GR and DK conceived the study and planned the research. GR performed and analysed most experiments with help from VLSK, WKY, JH, IN and AJ. SW and FL performed and analysed the crystallography experiments with help from YS. GR, VLSK and DK established the ZBTB48 knock-out mouse strain. GR, MMHK and DK analysed bone-marrow samples with help from MO. VK and VM analysed the omics datasets. GR and DK wrote the manuscript with input from all authors.

## DECLARATION OF INTERESTS

The authors declare that they have no competing interests.

## DATA AVAILABILITY

The atomic coordinates and structure factors for the ZBTB48 DNA binding domains in complex with its CIITA promoter binding site have been deposited to the Protein Data Bank (PDB) under the accession code 8JQ0. The mass spectrometry data have been deposited to the ProteomeXchange Consortium via the PRIDE partner repository^70^ with the dataset identifier PXD043134. The RNA-seq, ChIP-seq and FAIRE-seq data are available at the NCBI Gene Expression Omnibus (GEO)^71^ with the identifier GSE237959.

## SUPPLEMENTARY FIGURES

**Figure S1:**
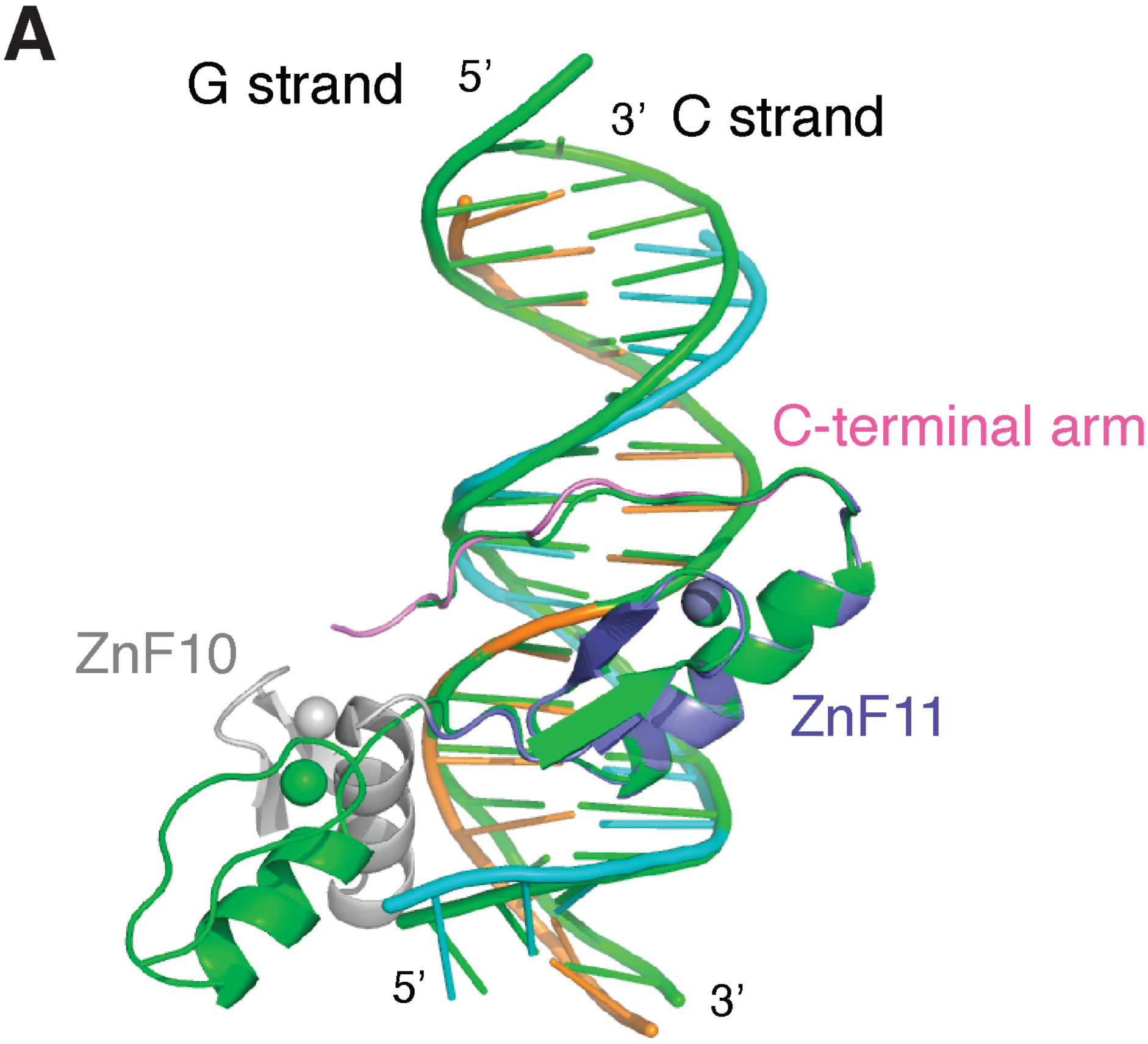
ZBTB48 binding to telomeric DNA vs. CIITA pIII are differentiated by ZnF10. (A) Superposition of the co-crystal structure of the essential ZBTB48 DNA-binding domains with telomeric DNA (bright green) and the e.2 binding site within CIITA pIII (colored as in Fig. 2B). Note how ZnF10 contributes to the interaction with the e.2 sequence while lies outside the DNA duplex in case of the telomeric DNA.

**Figure S2:**
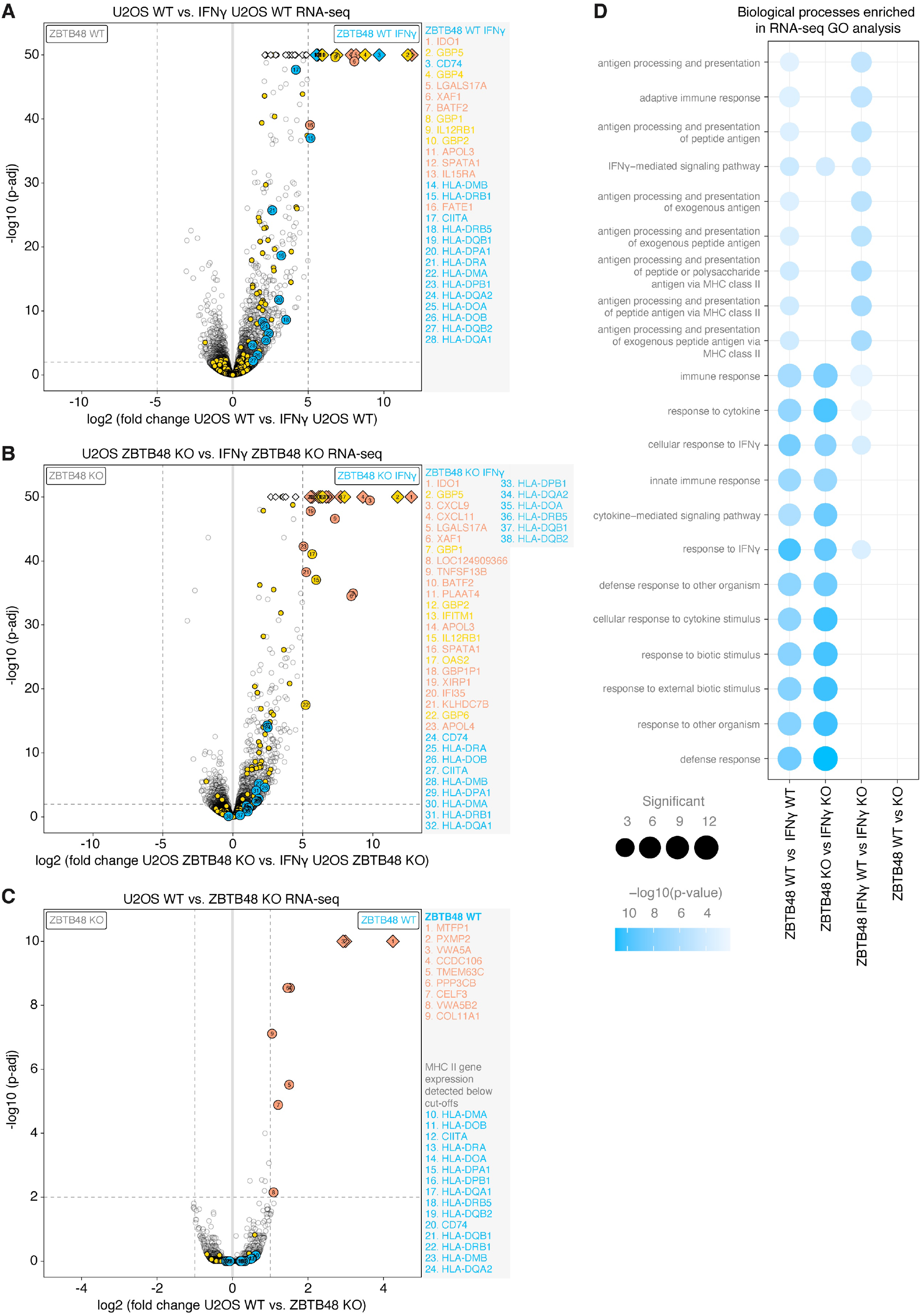
ZBTB48 exclusively affects the CIITA-MHC II expression program upon IFNγ stimulation at the mRNA level. (A) Differential expression analysis of the RNA-seq quantification comparing 5 U2OS WT clones +/- IFNγ treatment (250 U/ml IFNγ for 24 hours). Genes belonging to the CIITA-MHCII family are in blue and the rest of the IFNγ response genes (ISGs) in yellow. Additional proteins that expressed above a two-dimensinal cut-off of >32-fold enrichment (owing to the large number of strongly induced genes) and p<0.01 are in salmon. (B) Differential expression analysis of the RNA-seq quantification comparing 5 U2OS ZBTB48 KO clones +/- IFNγ treatment (250 U/ml IFNγ for 24 hours) depicted as in (A). (C) Differential expression analysis of the RNA-seq quantification comparing 5 U2OS WT and ZBTB48 KO clones (without IFNγ treatment). Genes are colored as in (A) and (B), but here a >2-fold enrichment cut-off was applied (as in Fig. 3D) given that baseline expression differences are compared. (D) GO term analysis for enriched proteins in pair-wise comparisons in Fig. 3D and Fig. S2A-C depicting significantly enriched GO terms. The size of the dots corresponds to the number of genes representing each term.

**Figure S3:**
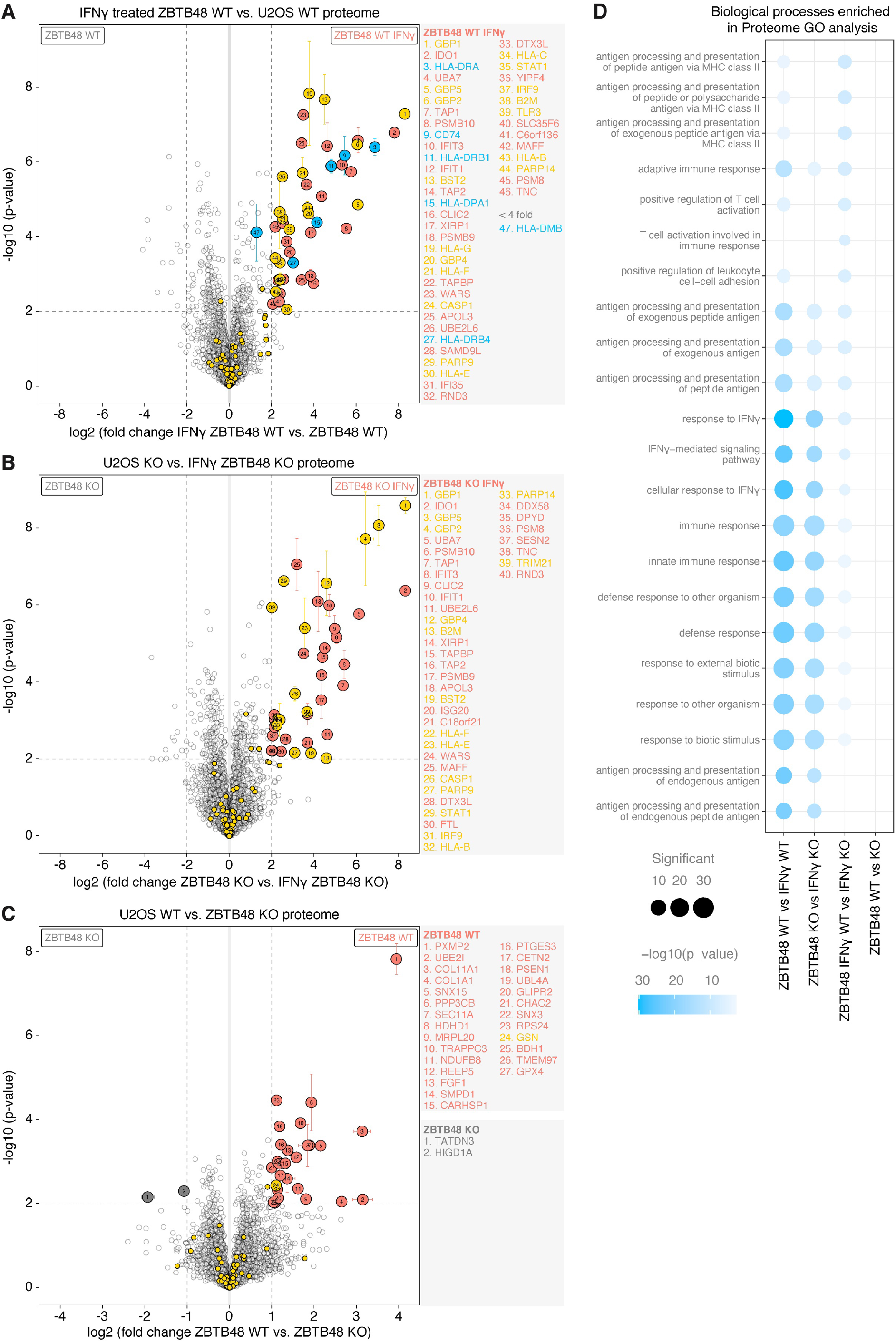
ZBTB48 exclusively affects the CIITA-MHC II expression program upon IFNγ stimulation at the protein level. (A) Differential expression analysis of label-free quantitative protein expression comparing 5 U2OS WT clones +/- IFNγ treatment (250 U/ml IFNγ for 48 hours). Genes belonging to the CIITA-MHCII family are in blue and the rest of the IFNγ response genes (ISGs) in yellow. Additional proteins that expressed above a two-dimensinal cut-off of >4-fold enrichment (owing to the large number of strongly induced genes) and p<0.01 are in salmon. (B) Differential expression analysis of the RNA-seq quantification comparing 5 U2OS ZBTB48 KO clones +/- IFNγ treatment (250 U/ml IFNγ for 48 hours) depicted as in (A). (C) Differential expression analysis of the RNA-seq quantification comparing 5 U2OS WT and ZBTB48 KO clones (without IFNγ treatment). Genes are colored as in (A) and (B), but here a >2-fold enrichment cut-off was applied (as in Fig. 3F). (D) GO term analysis for enriched proteins in pair-wise comparisons in Fig. 3F and Fig. S3A-C depicting significantly enriched GO terms. The size of the dots corresponds to the number of genes representing each term.

**Figure S4:**
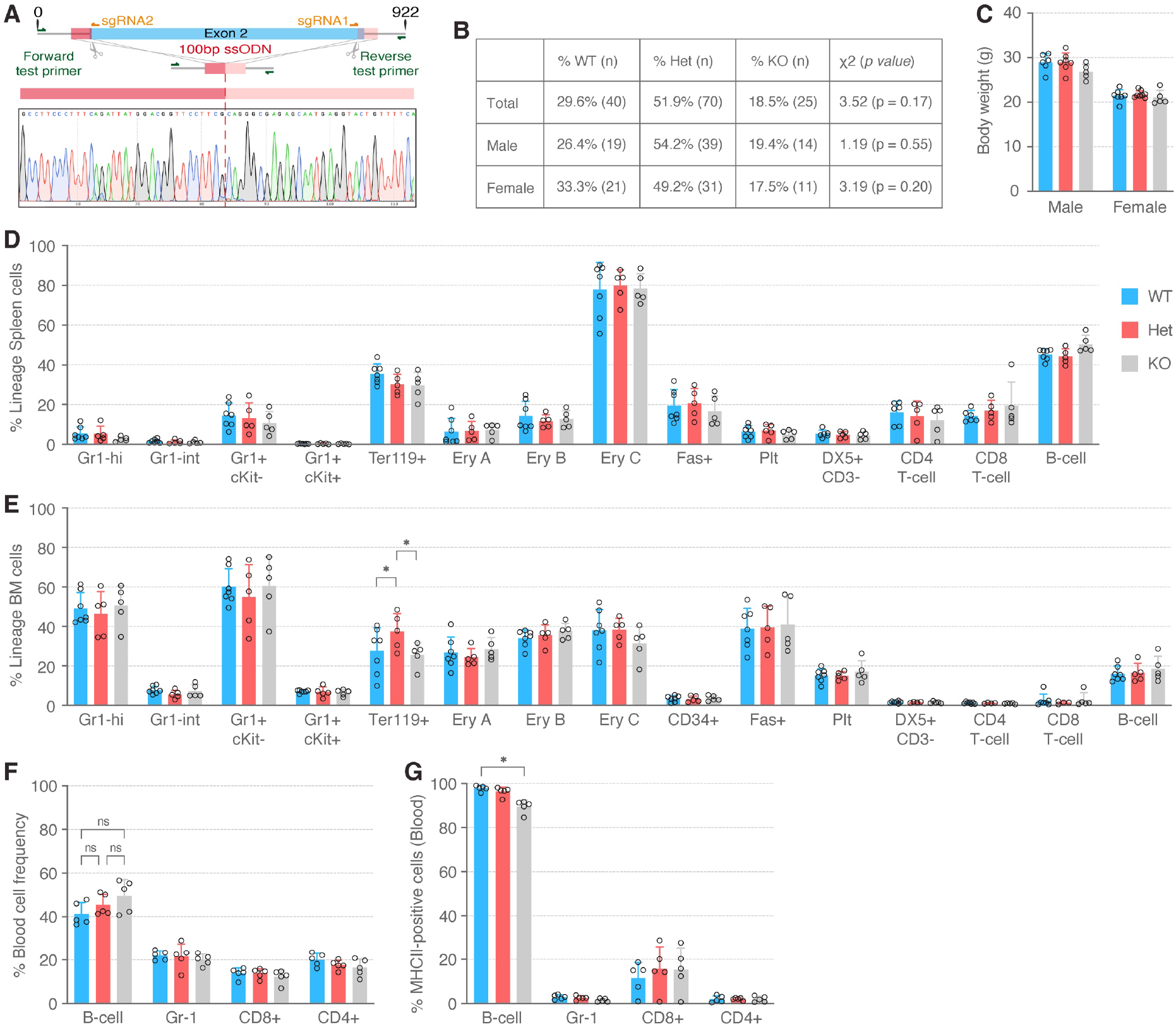
Loss of ZBTB48 does not affect abundance of blood cell populations. (A) Schematic of the sgRNA and ssODN design used for creation of the ZBTB48 KO strain (top) and Sanger sequencing verification of successful and precise deletion (bottom) in ZBTB48^-/-^ animals. (B) χ^2^-test comparing experimentally determined offspring frequency in crosses between ZBTB48^+/-^ animals. The genotype distribution follows Mendelian inheritance patterns for the total cohort (top), male (middle) and female (bottom) mice. (C) Body weight in g calculated separately for males and females across the three genotypes. Data represents mean ± SD. (male: WT n = 6, Het n = 7, KO n = 5; female: WT n = 7, Het n = 8, KO n = 5). These data were used to generate the ratio with the spleen weight in Fig. 4D. (D) Percentage of spleen lineage cells in WT (n = 7), Het (n = 5) and KO (n = 5) mice. (E) Percentage of bone marrow lineage cells in WT (n = 7), Het (n = 5) and KO (n = 5) mice. (F) Percentage of blood cell frequencies in WT (n = 7), Het (n = 5) and KO (n = 5) mice. (G) Percentage of MHCII-positive cells in the blood cell types from Fig. S4F. For (D-G) data represents mean ± SD. p-values were calculated by 2-way ANOVA with Sidak correction for multiple comparisons.

**Figure S5:**
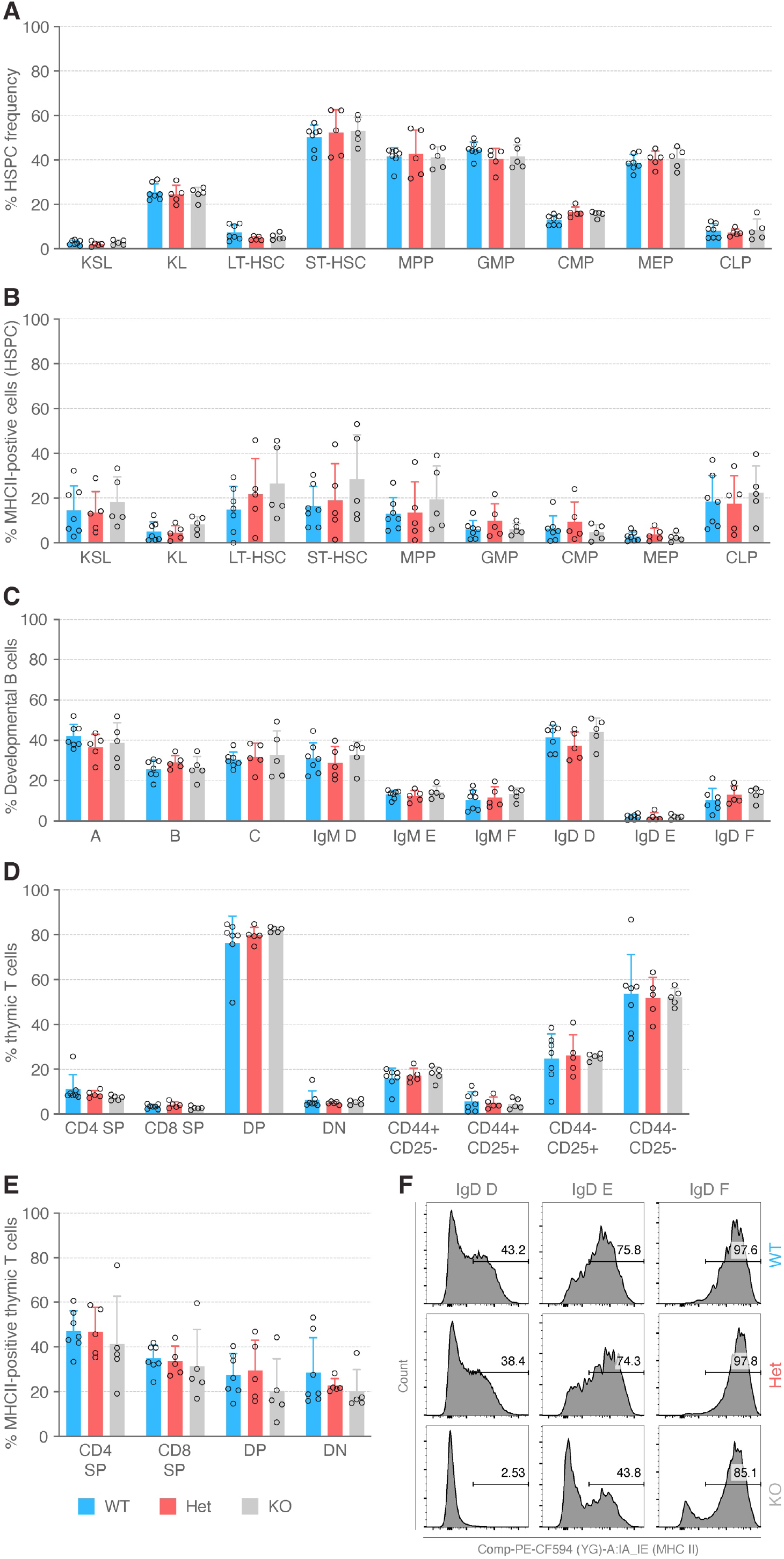
Loss of ZBTB48 does not affect abundance of hematopoietic stem cells, developmental B cells and thymic T cells. (A) Percentage of hematopoietic stem and progenitor cell (HSPC) types in WT (n = 7), Het (n = 5) and KO (n = 5) mice. (B) Percentage of MHCII-positive cells in HSPCs from Fig. S5A. (C) Percentage of development B cell types in WT (n = 7), Het (n = 5) and KO (n = 5) mice. (D) Percentage of thymic T cells in WT (n = 7), Het (n = 5) and KO (n = 5) mice. (E) B) Percentage of MHCII-positive cells in thymic T cells from Fig. S5D. For (A-E) data represents mean ± SD. p-values were calculated by 2-way ANOVA with Sidak correction for multiple comparisons. (F) Representative flow cytometry analysis of MHC II positive cells in D (pre-B), E (immature) and F (mature) subpopulations from WT, Het and KO littermates when gated by IgD corresponding to Fig. 4H-I.

**Figure S6:**
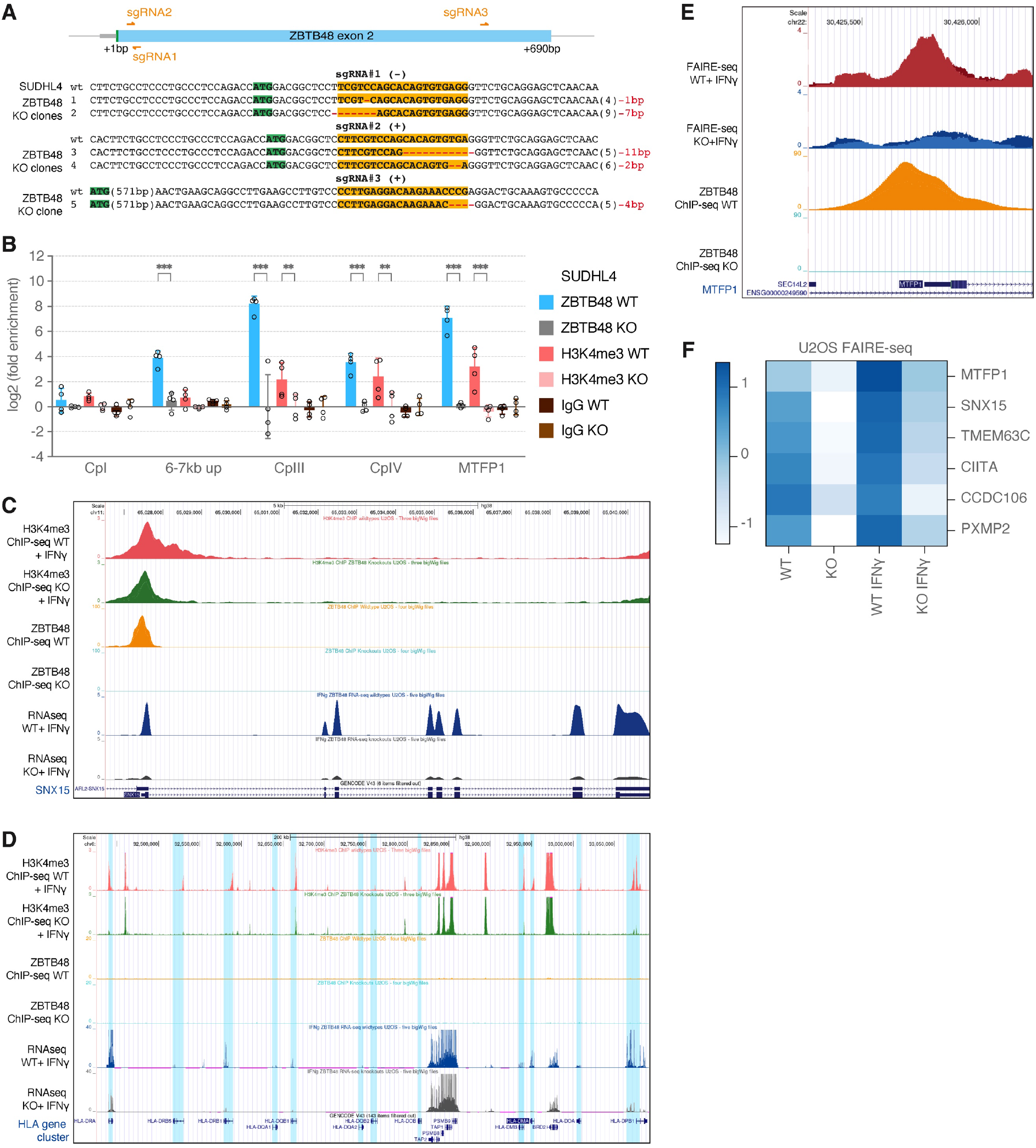
ZBTB48 also binds to CIITA pIII and regulates H3K4me3 levels in the constitutively expressing DLBCL cell line SUDHL4. (A) Schematic of sgRNA design for the generation of ZBTB48 KO clones in SUDHL4. 5 different KO clones were identified and genotyped as shown. The start codon is shown in green and the sgRNA binding sites are highlighted in bold and yellow. (B) ZBTB48 and H3K4me3 ChIP reactions from four independent SUDHL4 WT and ZBTB48 KO clones analysed by qPCR for CIITA pI, pII (6-7kb up), pIII, pIV. The MTFP1 promoter is used as a positive control. The data was normalised to IgG and gene desert region and fold-change was calculated relative to the average of WT clones. Data represents mean ± SD. p-values were calculated by 2-way ANOVA with Sidak correction for multiple comparisons (n = 4). (C) ChIP-seq tracks depicting the ZBTB48 at the SNX15 promoter in U2OS WT cells but not in the KO cells with concomitant loss of RNA expression as seen in the RNA-seq tracks. In contrast, the H3K4me3 peak is present at the SNX15 promoter in both U2OS WT and ZBTB48 KO clones. (D) ChIP-seq tracks as in (C) covering the MHC II gene cluster. The location of individual HLA genes is highlighted in light blue across tracks. (E) FAIRE-seq tracks depicting open chromatin at the MTFP1 promoter in two independent IFNγ-induced U2OS WT cells but not in the KO cells. The ZBTB48 ChIP-seq tracks highlight the ZBTB48 binding region within the MTFP1 promoter. (F) Heatmap of FAIRE-seq profiles depicting the relative chromatin accessibility at ZBTB48 bound promoter in U2OS WT vs ZBTB48 KO cells +/- IFNγ treatment (250 U/ml IFNγ for 48 hours). Intensity values are based on Z-scores of log2 rpkm values across samples.

## SUPPLEMENTARY TABLES

**Table S1.** Co-crystal structure data collection and refinement statistics.

|  | ZBTB48-CIITA DNA |
| --- | --- |
| Wavelength(Å) | 0.979 |
| Space group | $P4_32_12$ |
| Cell parameters |  |
| a, b, c (Å) | 126.56, 46.86, 79.05 |
| $\alpha$, $\beta$, $\gamma$ (°) | 90, 124.89, 90 |
| Resolution (Å) | 40.00-2.90 (2.95-2.90)<sup>a</sup> |
| $R_{\text{merge}}$ (%) | 11.7 (86.7) |
| $CC_{1/2}$ (%) | 98.9 (81.5) |
| $I/\sigma I$ | 14.6 (1.0) |
| Completeness (%) | 98.9 (91.4) |
| Average redundancy | 6.2 (5.0) |
| **Refinement** |  |
| No. reflections (overall) | 7083 |
| No. reflections (test set) | 330 |
| $R_{\text{work}}/R_{\text{free}}$(%) | 23.34/29.12 |
| Protein | 1172 |
| DNA | 1300 |
| ZN | 4 |
| B factors (Å<sup>2</sup>) |  |
| Protein | 46.44 |
| DNA | 69.02 |
| ZN | 30.58 |
| R.m.s. deviations |  |
| Bond lengths (Å) | 0.003 |
| Bond angles (°) | 0.473 |
| RAMPAGE ramachandran plot % residues |  |
| Favored | 92.42 |
| Allowed | 7.58 |
| Outliers | 0 |

**Table S2 | U2OS WT and ZBTB48 KO RNA-seq data (+/- IFNγ treatment)**

**Table S3 | U2OS WT and ZBTB48 KO proteome data (+/- IFNγ treatment)**

**Table S4 | U2OS WT and ZBTB48 KO (+ IFNγ treatment) H3K4me3 ChIP-seq data**

**Table S5 | U2OS WT and ZBTB48 KO (+ IFNγ treatment) FAIRE-seq data**

**Table S6 | Oligonucleotides used in this study**

**Table S7 | Antibodies used in this study**

## Notes

### Competing Interest Statement

The authors have declared no competing interest.

